# Deep learning localizes focal biology implicated in therapeutic resistance within routine histology

**DOI:** 10.64898/2026.05.05.723013

**Authors:** Tiago Goncalves, Dagoberto Pulido, Carmen M. Perrino, Thanpisit Lomphithak, Mason Cleveland, Adrian V. Dalca, Elizabeth Gerstner, Jason Hipp, Jay Patel, Bruce Rosen, Sahussapont Joseph Sirintrapun, Seth A. Wander, Anil Parwani, Gary Tozbikian, M. Khalid Khan Niazi, Jaime Cardoso, Jane Brock, Valentina Zanfagnin, Francesca Gazzaniga, A. John Iafrate, Keith T. Flaherty, Dennis C. Sgroi, John V. Guttag, Christopher P. Bridge, Albert E. Kim

**Affiliations:** INESC TEC, Faculdade de Engenharia, Universidade do Porto, Rua Dr. Roberto Frias, 4200-465 Porto, Portugal; Athinoula A. Martinos Center for Biomedical Imaging, Massachusetts General Hospital, Harvard Medical School; Boston, MA; Department of Pathology, Massachusetts General Hospital, Harvard Medical School; Boston, MA; Computer Science and Artificial Intelligence Laboratory (CSAIL), Department of Computer Science, Massachusetts Institute of Technology; Cambridge, MA; Massachusetts General Hospital Cancer Center, Harvard Medical School; Boston, MA; Department of Pathology, The Ohio State University; Columbus, OH; American Society of Clinical Pathology; Chicago, IL

**Author notes:** **Corresponding authors** Albert E. Kim, M.D. Simches Research Building, 185 Cambridge Street, Office 3.828, Boston, MA 02114, Christopher P. Bridge, D.Phil. 149 13^th^ Street, Office 2303, Boston, MA 02129. co-senior author (Christopher P. Bridge, Albert E. Kim).

## Abstract

Precision oncology lacks scalable methods to identify the mechanisms that mediate therapeutic resistance for individual patients. Resistance often arises from focal cellular niches that are obscured by bulk profiling and costly to resolve with multi-omics. Here, we show that deep learning (DL), applied to routine histology, can localize focal tissue regions enriched for therapeutically relevant biology. Using 3111 breast cancer H&E slides with matched bulk transcriptomics, we trained weakly-supervised DL models to infer activities of immune, metabolic, and tumor-intrinsic phenotypes implicated in therapeutic resistance (AUROC>0.80; PCC>0.64). Accurate inference of these phenotypes should identify tissue regions enriched for the corresponding biological signal. Therefore, we validated phenotype inference and spatial localization with complementary analyses. Tissue-matched multiplexed immunofluorescence showed concordance between inferred immune states and corresponding cell fractions (p=0.006-0.106). Across multi-institutional cohorts, model-derived phenotypes recovered expected relationships with therapeutic outcomes (p<0.045). Finally, in a blinded evaluation, pathologists confirmed that model-derived high-attention regions were enriched for phenotype-specific morphology (p<2.408*10^-5^). Because evaluated phenotypes represent diverse mechanisms of resistance across therapeutic modalities, these findings provide a foundation for *resistance-directed localization* using therapeutic outcomes as supervision. By directing deep profiling toward model-prioritized regions, this framework could enable scalable nomination of candidate mediators of resistance for subsequent functional validation across real-world patient populations.

**One sentence summary:** Weakly supervised deep learning localizes focal tissue regions enriched for therapeutically relevant biology in routine histology, thus offering a scalable strategy to study therapeutic resistance across large patient populations.

## INTRODUCTION

A fundamental goal of precision oncology is to uncover the patient-specific biological mechanisms that mediate therapeutic resistance. Despite remarkable advances in cancer genomics, our understanding of resistance biology remains bottlenecked by the scale and spatial resolution of current tools. Emerging evidence suggests that therapeutic efficacy across cancer types can be disproportionately shaped by focal niches of cancer cells or sparsely represented populations that may be overlooked by bulk or spatially agnostic assays^1–5^. For example, small subpopulations of immune cells can influence response to checkpoint inhibition. For oncogene-directed therapies, rare malignant subclones or immunosuppressive cell populations can persist during treatment and contribute to resistance. Yet defining these cellular niches typically requires spatial profiling, which remain resource-intensive and largely restricted to small research cohorts. In contrast, companion diagnostics used in routine care provide a limited and spatially unresolved view of tumor biology. Consequently, mechanistic insights are largely derived from small patient cohorts and may not capture the diversity of therapeutically relevant biology within real-world patient populations. This gap highlights a pressing need for scalable methods to identify focal cellular niches that mediate therapeutic efficacy.

Modern cancer centers have amassed large repositories of patient-derived clinical data, while advances in deep learning (DL) and high-performance computing enable the extraction of latent biology from routine clinical data, such as digital pathology^6–9^. Together, these developments create an opportunity to study therapeutic resistance through a new lens: using pathology generated during clinical care to characterize patient-specific biology. Our approach rests on the premise that routine H&E slides encode critical elements of patient-specific tumor microenvironment (TME) biology, such as tissue architecture and the spatial organization of diverse cell populations. However, the scale of whole-slide images (WSIs) preclude reproducible, quantitative integration of these features through manual review alone. DL provides a means to analyze this information across entire WSIs and large patient populations.

Previous computational pathology studies have shown that DL can predict molecular states and spatial transcriptomic (ST) profiles from routine histology^10–12^. ST prediction generally seeks to reconstruct predefined molecular measurements throughout the tissue. Our complementary strategy instead uses patient-level biological states to prioritize focal regions most strongly associated with the corresponding biology for focused molecular investigation. We selected breast cancer, the most frequently diagnosed cancer worldwide, as our initial application because its biological heterogeneity is incompletely captured by standard profiling methods (e.g., estrogen and progesterone receptor immunohistochemistry [IHC], HER-2 IHC/FISH). Using patient-matched bulk RNA sequencing and H&E WSIs, we trained DL models to quantify TME programs implicated in therapeutic resistance and to prioritize the focal tissue regions associated with each program. We evaluated the validity of this approach using tissue-matched multiplexed immunofluorescence and therapeutic outcome associations across multi-institutional cohorts. We then directly evaluated spatial localization through a blinded pathologist study that tested whether model-prioritized regions were enriched for phenotype-specific morphology.

Together, these analyses showed that weakly supervised models could infer therapeutically relevant TME phenotypes while localizing focal regions enriched for phenotype-specific biology. Because the evaluated TME phenotypes represent diverse resistance mechanisms across cancer types and treatment modalities, our findings provide a foundation for applying DL-based localization to the study of therapeutic resistance. By using therapeutic outcomes as supervision, this framework could prioritize focal tissue regions for mechanistic investigation.

## RESULTS

### Deep learning infers TME phenotypic biology from H&E WSIs

Using tissue-matched breast cancer H&E WSIs (n = 3111 WSIs; 1098 patients) and bulk RNA-sequencing data, we trained weakly supervised DL models to quantify activity of TME programs implicated in therapeutic resistance directly from routine histology. Ground truth functional states for ten TME phenotypes spanning tumor-intrinsic, immune, and metabolic programs were defined for each patient using single-sample gene set enrichment analysis (ssGSEA). On the held-out test set, our DL models accurately quantified these systems-level phenotypes (**Figure 2**). Performance was similar across biological systems, including immune (area under receiver operating characteristic curve [AUROC]: 0.861 {95% confidence interval: 0.805-0.907}; Pearson’s correlation coefficient [PCC]: 0.789 {0.731-0.840}, cancer cell-intrinsic (AUROC: 0.861 [0.811-0.911]; PCC: 0.744 [0.671-0.802]), and metabolic phenotypes (AUROC: 0.843 [0.791-0.890]; PCC: 0.741 [0.664-0.812]). Subgroup analysis across our test set revealed receptor profile-specific differences in performance, although this observation is tempered by the small number of patients in the HR+/HER2-cohort (**Supplemental Table 1**). These models also outperformed previously reported histology-based approaches for predicting individual gene or protein expression^13,14^, supporting the use of systems-level phenotypes as supervision.

**Figure 1:**
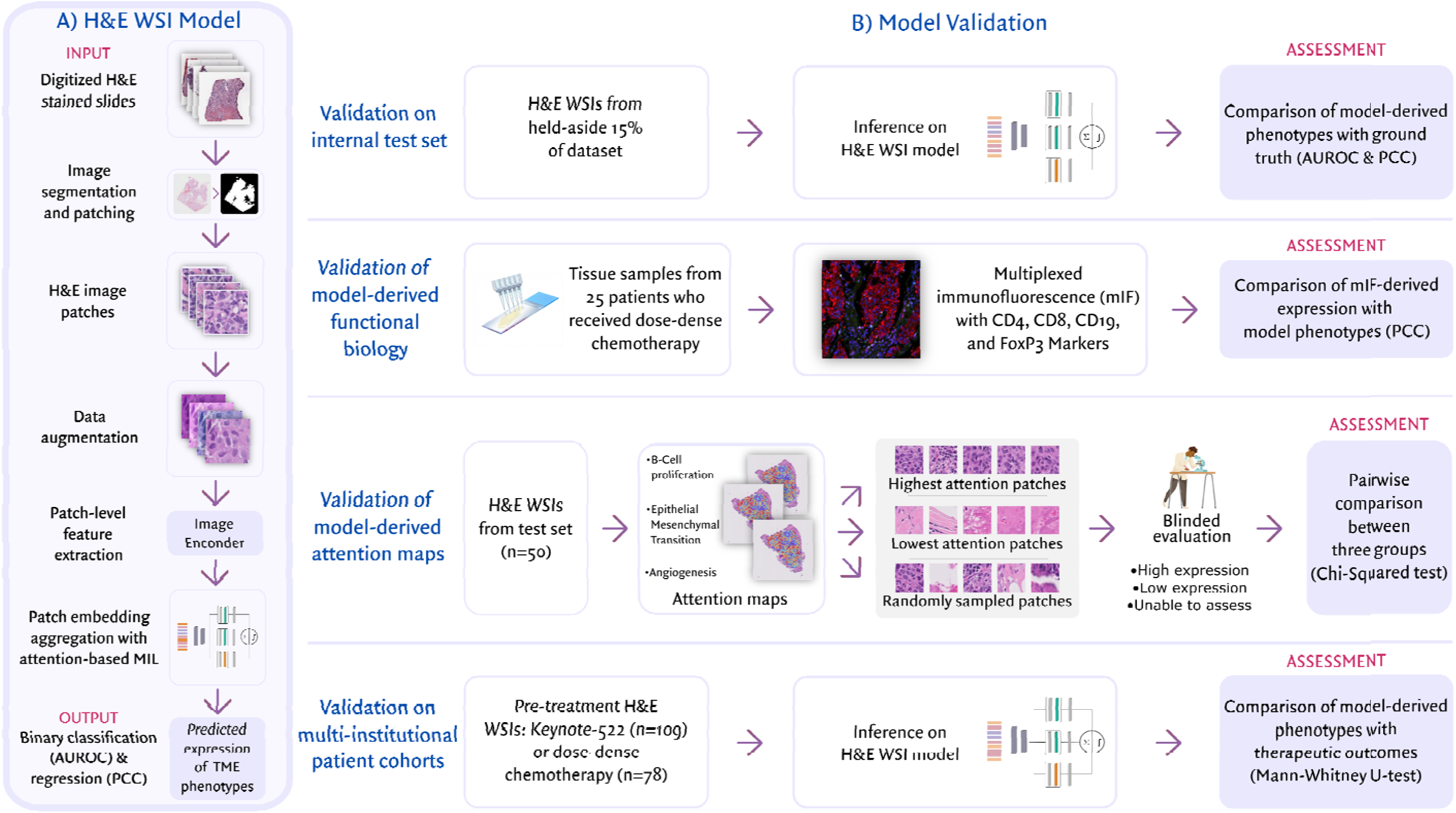
Study design, model training, and validation strategy. A) **H&E WSI model development.** The ground truth for each H&E WSI was defined by tissue-matched bulk RNA-sequencing data. Following tissue segmentation and quality control, non-overlapping patches of 256 x 256 pixels were extracted from the tissue regions of the WSIs. Color augmentation and geometric transforms were applied to these image patches to mimic common sources of variability of H&E WSIs. Patch level features were extracted via an image encoder, and attention-weighted aggregation was employed to generate a single slide-level feature vector. This vector was passed to task-specific prediction heads using an attention-based multiple instance learning (MIL) framework. Binary classification and regression models were trained to quantify each TME phenotypic activity, with performance evaluated using area under the receiver operating characteristic (AUROC) and Pearson’s correlation coefficient (PCC) metrics, respectively. B) **Model validation.** The ability of our model to accurately resolve systems-level TME phenotypes with *weak supervision* was validated using four distinct methods. • Internal validation: Model performance was assessed on the held-aside test set, where predictions were compared directly with ground-truth transcriptomic-derived phenotypic activity. • Validation of functional biology: The model’s ability to disentangle different immune phenotypes was assessed using tissue matched multiplexed IF from our multi-institutional cohort (n = 25). PCCs were calculated to evaluate the association between model-derived phenotypes and corresponding mIF-defined cellular fractions (CD19/20/45+ B cells; CD4/GZMB+ and CD8/GZMB+ cytotoxic T cells; CD4/FoxP3+ regulatory T cells) • Validation of attention maps: A blinded reader study was conducted to assess the biological fidelity of attention maps. For fifty randomly selected H&E WSIs within our test set, two pathologists blinded to the model output evaluated three groups of patches (e.g., those with the highest attention scores, the lowest attention scores, and randomly sampled patches) to determine if observed histologic features were consistent with known phenotypic activity. • Validation on multi-institutional patient cohorts: Model generalizability and clinical utility were evaluated on multi-institutional cohorts (n = 199) treated with either neoadjuvant pembrolizumab and chemotherapy (Keynote-522) or adjuvant dose-dense chemotherapy. Clinical outcomes were correlated with model-derived immune and cell cycling phenotypes, in accordance with established Omics correlates of therapeutic efficacy.

**Figure 2:**
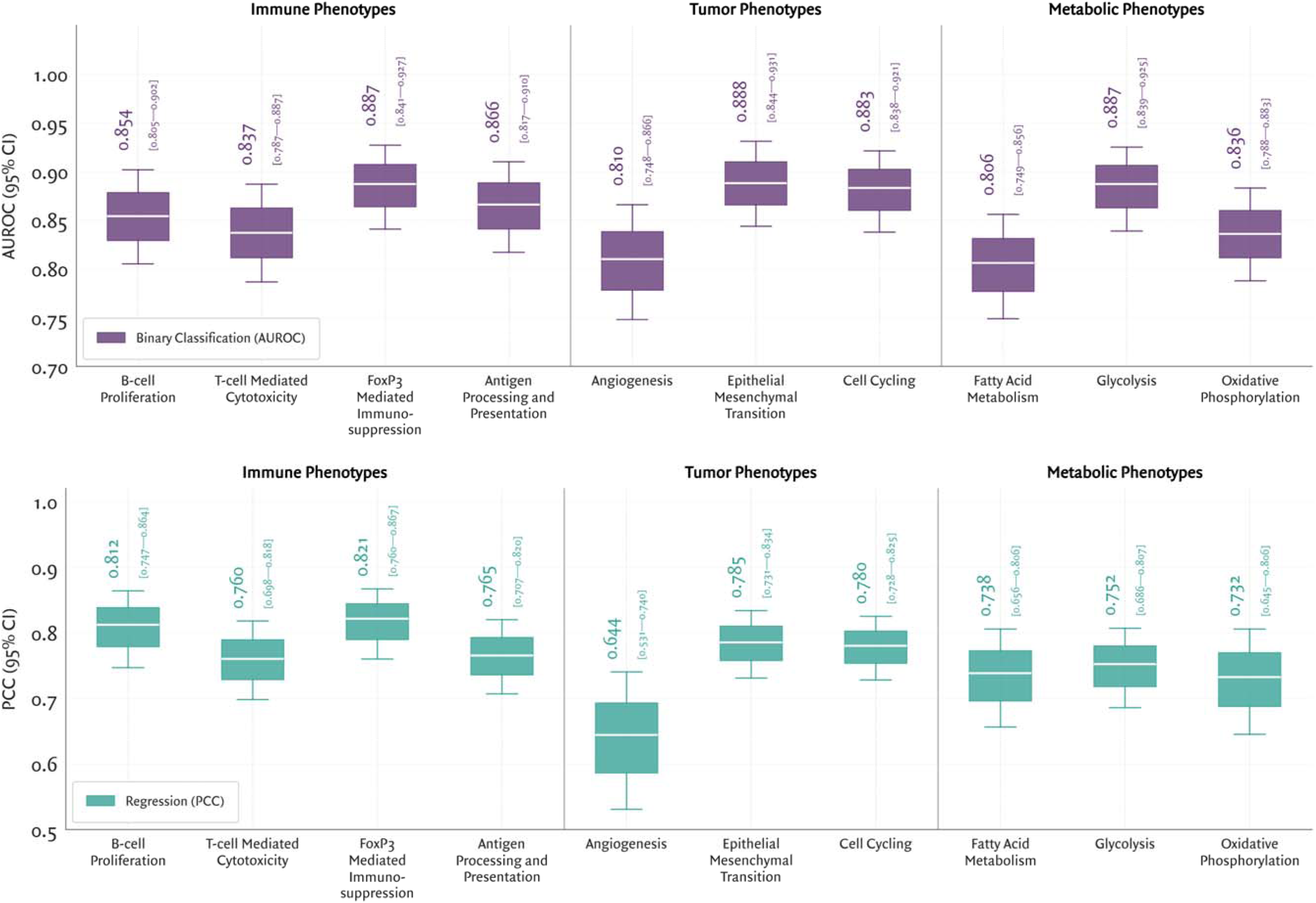
Deep learning infers therapeutically relevant systems level tumor microenvironment biology from routine H&E WSIs. Model performance is shown for ten pre-defined TME phenotypes, spanning immune, tumor, and metabolic phenotypes, relevant for therapeutic resistance across the pan-cancer landscape. For each phenotype, model performance is reported for binary classification (AUROC; purple) and regression (PCC, teal). Box-and-whisker plots display the median and 95% confidence intervals estimated via bootstrap resampling across the held-aside test set. Across all tasks, models achieved AUROC > 0.80 for binary classification and PCC > 0.64 for regression, suggesting that tissue architecture and spatial distribution of cell populations within H&E slides encode system levels TME biology.

Models using tabular clinico-genomic data commonly used for patient stratification performed worse than WSI-only models (**Supplemental Table 2**). These tabular models showed relatively stronger performance (AUROC 0.70-0.80) for cell cycling and glycolysis phenotypes, likely reflecting the higher expression of these programs in certain molecular subtypes. For example, TNBCs frequently exhibit elevated aerobic glycolysis, which supports rapid proliferation and immunosuppression, compared to hormone receptor-positive tumors^15,16^. Consistent with these trends, 75.62% and 77.50% of the TNBC tumors in our dataset exhibited up-regulation of cell cycling and glycolytic activity, respectively. These findings indicate that routine histology contains biological information not captured by conventional variables for patient stratification.

We evaluated several strategies to optimize model performance. First, we compared open-source image encoders to identify the optimal feature representation for TME phenotyping. Since the CONCH vision-language foundation model outperformed PLIP and ResNET-50 (**Supplemental Table 3**), all subsequent experiments in this work used CONCH-derived features. Next, to further improve model generalization, we implemented label-preserving data augmentation strategies, including patch-level geometric transformations and color augmentations. These augmentations improved performance for most phenotypes compared to models trained without them (**Supplemental Figure 1**). Finally, we compared performance of attention-based MIL with a vision transformer-based architecture (TransMIL)^17^. Performance was similar between these architectures, suggesting that modeling long-range spatial dependencies did not improve phenotype inference at the current dataset size.

### Multimodal integration of H&E WSIs and clinico-genomic data does not improve performance

We next evaluated whether the integration of routinely acquired clinico-genomic metadata (e.g., age, tumor grade, stage, receptor profile) with H&E WSIs improves inference of TME phenotypes. We evaluated two multi-modal fusion strategies: (i) intermediate cross-attention fusion, adapted from the SurvPath architecture^18^ to capture inter-modality interactions, and (ii) late fusion, in which modality-specific predictions were combined with a multi-layer perceptron. Performance of these multimodal models was benchmarked against the H&E WSI model on the held-out test set.

Across nearly all phenotypes, multimodal models yielded no measurable improvement over the H&E WSI baseline. In several instances, particularly for the cross-attention models, inclusion of clinical features reduced model performance (**Supplemental Figure 2**). Feature ablation showed that predictive signal in the clinico-genomic models was driven primarily by molecular data (e.g., ER, PR, HER-2 status), with negligible contributions from demographics or tumor stage (**Supplemental Table 4**). Moreover, the addition of any clinico-genomic feature category to the multimodal models provided no measurable improvement (**Supplemental Table 5**). Together, these results suggest that evaluated clinico-genomic features provided little additional information beyond that encoded in routine histology.

### Model-derived immune phenotypes align with tissue-matched protein expression

Our central premise is that accurate inference of TME phenotypes requires the model to correctly detect the corresponding biology encoded in the H&E slides. First, we evaluated whether inferred model phenotypes reflected orthogonal protein-level measurements by performing tissue-matched mIF on 25 tissue samples from MGH. For each sample, the matched H&E WSI was processed through our final model. The mIF panel defined B cells, cytotoxic T cells, and regulatory T cells (Tregs) by employing antibodies against CD4, CD8, CD19, CD20, CD45, granzyme B (GZMB), and FoxP3. Although these markers quantify canonical cell populations related to our model’s immune phenotypes, the transcriptomic signatures used for model supervision encompass broader sets of functional genes. We therefore expected correspondence, but not equivalence, between model-derived phenotypes and mIF-derived cell fractions. PCCs were calculated to assess these associations.

Model-derived T-cell cytotoxicity expression was significantly associated with the fraction of cytotoxic T cells (PCC: 0.534; p = 0.006; **Table 1**), supporting the model’s ability to infer state-specific immune function from H&E WSIs (**Supplemental Figure 3**). Similarly, FoxP3-mediated immunosuppression was significantly associated with the fraction of Tregs (PCC: 0.437; p = 0.029; **Table 1, Supplemental Figure 4**). Review of discrepant cases identified FoxP3 expression in other cell populations (e.g., CD11b+ myeloid cells and PanCK+ tumor cells; **Supplemental Figure 5**). These observations suggest that the model-derived phenotype may capture a broader FoxP3-associated immunosuppressive state rather than Treg abundance alone.

**Table 1:** Model-derived phenotypic activity is linked with multiplexed immunofluorescence-defined fractions of canonical cell populations. . Cellular fractions were derived from 25 tissue-matched tumor samples within the ddACT cohort. mIF gating strategies defined B cells as CD19/CD20/CD45-positive; cytotoxic T cells as CD4/GZMB-positive and CD8/GZMB-positive; and regulatory T cells as CD4/FoxP3-positive. Fractions were calculated relative to the total cell count per sample. Statistically significant positive correlations were observed between model-derived T-cell cytotoxicity and mIF-defined cytotoxic T cell fractions (*r* = 0.534; p = 0.006), as well as model-derived FoxP3-mediated immunosuppression and fraction of Tregs (*r* = 0.437; p = 0.029). B-cell proliferation demonstrated a positive trend with the mIF-defined B-cell fraction (*r* = 0.331; p = 0.106), perhaps reflecting the functional divergence between cellular density of B-cells and transcriptomic-based proliferative activity.

| Sample ID | B-cell proliferation |  | T-cell cytotoxicity |  | FoxP3-mediated immunosuppression |  |
| --- | --- | --- | --- | --- | --- | --- |
|  | Fraction of B cells (%) | Model-derived quantitative phenotypic activity | Fraction of cytotoxic T cells (%) | Model-derived quantitative phenotypic activity | Fraction of regulatory T cells (%) | Model-derived quantitative phenotypic activity |
| TN04 | 1.0143 | 0.9109 | 12.0239 | 2.1544 | 6.4866 | 0.9188 |
| TN12 | 0.0883 | 0.1827 | 2.0002 | 0.7356 | 0.7282 | 0.0488 |
| TN19 | 0.4905 | 0.0379 | 3.9243 | 0.3316 | 2.2547 | -0.1887 |
| TN21 | 0.0508 | -0.3256 | 3.4961 | -0.2867 | 0.6418 | 0.0493 |
| TN27 | 0.0311 | -0.1267 | 2.3280 | -0.0431 | 3.4717 | -0.1694 |
| TN30 | 0.0024 | -0.1066 | 0.9036 | 0.3811 | 0.8560 | 0.1063 |
| TN52 | 0.3755 | 0.5489 | 5.7815 | 1.5972 | 1.4326 | 0.4705 |
| TN53 | 8.9977 | 1.2606 | 12.7830 | 2.1239 | 16.5946 | 1.5677 |
| TN54 | 0.7880 | -0.0644 | 7.2795 | 1.9053 | 1.8355 | 0.4142 |
| TN57 | 0.0652 | 0.3018 | 4.5408 | 2.3541 | 1.3257 | 1.2354 |
| TN63 | 1.7561 | 1.1876 | 5.8682 | 2.3274 | 3.8394 | 1.0402 |
| TN64 | 0.0496 | 1.1784 | 2.4034 | 2.1693 | 1.6169 | 1.1857 |
| TN66 | 0.2012 | 1.4543 | 0.6626 | 1.5997 | 6.7022 | 0.2734 |
| TN68 | 0.1140 | 0.4629 | 1.7861 | 1.6275 | 0.3706 | 0.6477 |
| TN69 | 0.0236 | 1.0559 | 0.9601 | 1.7683 | 1.1466 | 1.0364 |
| TN74 | 0.0028 | 0.8243 | 0.7056 | 0.9206 | 4.0023 | 0.9460 |
| TN75 | 1.5887 | 0.7639 | 11.9110 | 1.9903 | 4.4656 | 0.7061 |
| TN80 | 1.6593 | 0.5888 | 7.1477 | 1.8002 | 2.4287 | 0.7171 |
| TN81 | 25.9727 | 1.0710 | 5.8876 | 1.8973 | 1.5193 | 1.2011 |
| TN93 | 0.1006 | 0.0911 | 3.9186 | 1.6058 | 0.5904 | 0.4003 |
| TN95 | 0.0000 | -0.3395 | 1.0876 | 1.5165 | 0.4124 | 0.1007 |
| TN97 | 0.1181 | 0.9587 | 8.8162 | 2.2929 | 4.0073 | 0.9501 |
| TN101 | 3.4829 | 1.2681 | 11.2163 | 2.1634 | 10.2394 | 1.1150 |
| TN107 | 0.0343 | 0.2184 | 0.6575 | 0.2063 | 4.6852 | 0.0785 |
| TN109 | 1.0784 | 0.5645 | 2.1096 | 1.8215 | 1.1409 | 0.7320 |

|  | Pearson’s correlation coefficient (r) | Statistical Significance |
| --- | --- | --- |
| *B-cell proliferation* | 0.331 | p = 0.106 |
| *T-cell cytotoxicity* | 0.534 | p = 0.006 |
| *FoxP3-mediated immunosuppression* | 0.437 | p = 0.029 |

Model-derived B-cell proliferation showed a positive but non-significant trend with mIF-derived B-cell fractions (PCC: 0.331; p = 0.106; **Table 1, Supplemental Figure 6**). This weaker association may reflect the distinction between total B-cell abundance, measured using CD19, CD20, and CD45, and the proliferative activity represented by the 98-gene transcriptomic signature used for model supervision. Overall, model-derived immune phenotypes aligned with corresponding mIF measurements. These results provide orthogonal protein-level evidence that model-derived immune phenotypes reflect corresponding TME biology. We next evaluated whether model inference recovered expected relationships with therapeutic outcomes across independent, multi-institutional patient cohorts.

### Model-derived phenotypes recover expected associations with therapeutic efficacy across multi-institutional cohorts

Next, we assessed validity of model-derived phenotypic signatures by assessing their therapeutic relevance in 188 patients treated at Massachusetts General Hospital (MGH), Brigham & Women’s Hospital (BWH), and Ohio State University. Whereas many DL models predict clinical outcomes directly, our framework was trained to recover TME phenotypes with established links to therapeutic resistance. We reasoned that accurate phenotype inference should recover expected relationships between these phenotypes and treatment efficacy without treatment-specific supervision. We evaluated this premise in two settings in which predictive biomarkers remain limited: (i) the neoadjuvant KEYNOTE-522 regimen (pembrolizumab with chemotherapy, followed by definitive surgery^19^) in stage II/III TNBC, and (ii) adjuvant dose-dense anthracycline and taxane-based chemotherapy (ddACT) in high-risk, early-stage TNBC.

For the KEYNOTE-522 cohort, we curated pre-treatment H&E WSIs from core biopsies of 109 TNBC patients who received neoadjuvant pembrolizumab with chemotherapy. Outcomes were dichotomized as pathologic complete response (pCR) or residual disease at time of definitive surgery. Given established associations between immunogenic tumor profiles and checkpoint inhibitor (ICI) efficacy^20,21^, analyses were restricted *a priori* to model-derived immune phenotypes. Consistent with these data, tumors achieving pCR showed higher model-derived activity for B-cell proliferation (p = 0.017), T-cell cytotoxicity (p = 0.020), and antigen processing and presentation (p = 0.045). Conversely, tumors from patients with residual disease exhibited higher model-derived FoxP3-mediated immunosuppression activity (**Table 3A**; p = 0.004). Thus, model-derived immune phenotypes recovered expected relationships with response to the KEYNOTE-522 regimen using only the pre-treatment core biopsy WSI.

For the adjuvant ddACT cohort, we curated pre-treatment H&E WSIs from 79 high-risk, early-stage TNBC patients with a minimum of seven years of follow-up. Outcomes were dichotomized as remission or relapse, defined by local recurrence or metastatic progression. Based on reported associations between chemotherapy efficacy with immune activation and tumor proliferative capacity^22–24^, analyses were restricted *a priori* to immune and cell cycling phenotypes. Although several associations trended towards the expected biological direction, they did not reach statistical significance (**Table 3B**). Differences between the two cohorts may reflect the distinct clinical endpoints and therapeutic settings. pCR provides a standardized assessment at a defined time point, whereas disease recurrence is a longitudinal outcome influenced by the time interval between treatment and relapse. The longer time interval may also reduce the association between the biology within a pre-treatment biopsy and the eventual clinical outcome.

Together, the multi-institutional analyses showed that model-derived phenotypes recovered expected phenotype-outcome relationships, providing additional evidence that the models inferred therapeutically relevant biology. Together with the mIF analyses, these findings support the premise that model attention prioritizes focal regions enriched for the relevant biological signal. We next tested this premise by evaluating whether pathologists could identify phenotype-specific morphology within high-attention regions.

### Attention localizes tissue regions enriched with phenotype-specific morphology

To determine whether our model’s predictions were driven by phenotype-relevant features rather than spurious correlates^25,26^, we examined the attention weights assigned to individual image patches. Each weight represented a patch’s contribution to the final WSI-level phenotype score. By projecting these weights onto the corresponding WSIs, we generated attention maps for the test set to evaluate whether high-attention regions contained recognizable histologic hallmarks. Qualitative review showed that attention maps highlighted spatially localized features consistent with the corresponding phenotype, such as dense lymphocytic aggregates for T-cell cytotoxicity and hypercellular tumor regions for cell cycling (**Figure 3**). A given image patch often received different attention scores across different phenotypes, indicating that the models prioritized regions in a task-specific manner within the same WSI. In contrast, low-attention regions exhibited tissue morphologies unrelated to the phenotype of interest.

**Figure 3:**
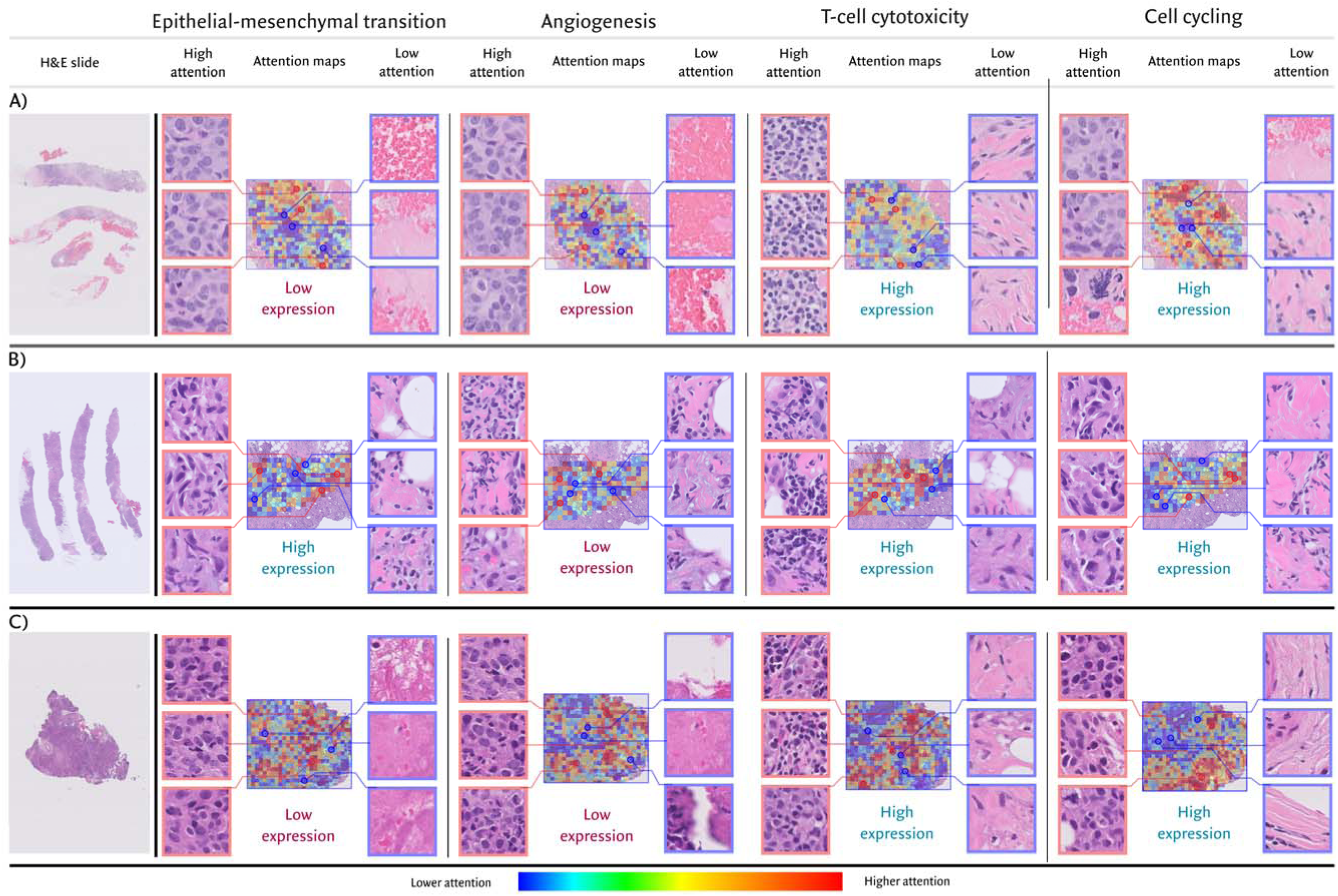
Weakly supervised attention mechanisms localize phenotype-specific TME biology within H&E WSIs. (A-C) Three representative cases illustrate the spatial localization of four TME functional programs derived from weak supervision on slide-level labels. For each case, an H&E WSI thumbnail (left) is presented alongside phenotype-specific attention heatmaps, and the top three high- and low-attention patches for each phenotype. The slide-level phenotype expression score, derived from tissue-matched bulk transcriptomics and used as ground truth, is labeled directly on each map in Red (Low expression) or Blue (high expression) text. The heatmap color scale (bottom) denotes patch-level attention weights from low (blue) to high (red). Across phenotypes, high-attention regions consistently correspond to visually explicit histologic structures consistent with the TME program of interest, such as hypercellular tumor regions for tumor-intrinsic phenotypes and lymphocyte-rich aggregates for immune phenotypes. High-attention regions additionally demonstrate distinct morphology, such as epithelioid cells for low expression of EMT and spindled, mesenchymal-like cells for high EMT expression. In contrast, low-attention regions were correctly assigned to morphologically uninformative areas, including adipose tissue, acellular eosinophilic material, or stroma. Together, these visualizations illustrate that attention scores prioritize tissue regions most informative for phenotype inference. Notably, the same image patch often received different, phenotype-specific attention scores, underscoring the model’s ability to spatially localize and assess different TME programs within the same WSI.

To quantitatively evaluate the biological fidelity of attention maps, we conducted a blinded reader study. Three phenotypes with recognizable morphologic correlates were evaluated: B-cell proliferation (lymphocytic infiltrates), epithelial-mesenchymal transition (invasive tumor cells and stromal fibroblasts), and angiogenesis (blood vessels with tumor cells). For each phenotype, 50 WSIs from the test set were selected. For each WSI, we generated three non-overlapping sets of five patches: the five patches with the highest attention scores, the five patches with the lowest attention scores, and five randomly sampled patches. Two board-certified pathologists (C.M.P.; J.A.H.) independently reviewed each patch set and predicted the corresponding slide-level phenotype as up-regulated, down-regulated, or unable to assess. Readers were blinded to the slide-level phenotype label, model prediction, and attention category. Patch sets were presented to the reader in random order. Reader performance was calculated separately for each pathologist and attention category.

Across all tasks, pathologist assessments based on high-attention patches consistently outperformed those using randomly sampled and low attention patches (**Table 2A**). For instance, high-attention patches reached a classification accuracy of up to 82% for B-cell proliferation, whereas low-attention patches were frequently determined to be “unable to assess” owing to irrelevant tissue morphology. Pairwise chi-squared testing confirmed that high-attention patches yielded significantly better reader performance than either comparator group (**Table 2B**; all p-values < 2.408 × 10^-5^). Together, these results provide direct evidence that model-prioritized tissue regions were enriched for phenotype-specific biology.

**Table 2:**
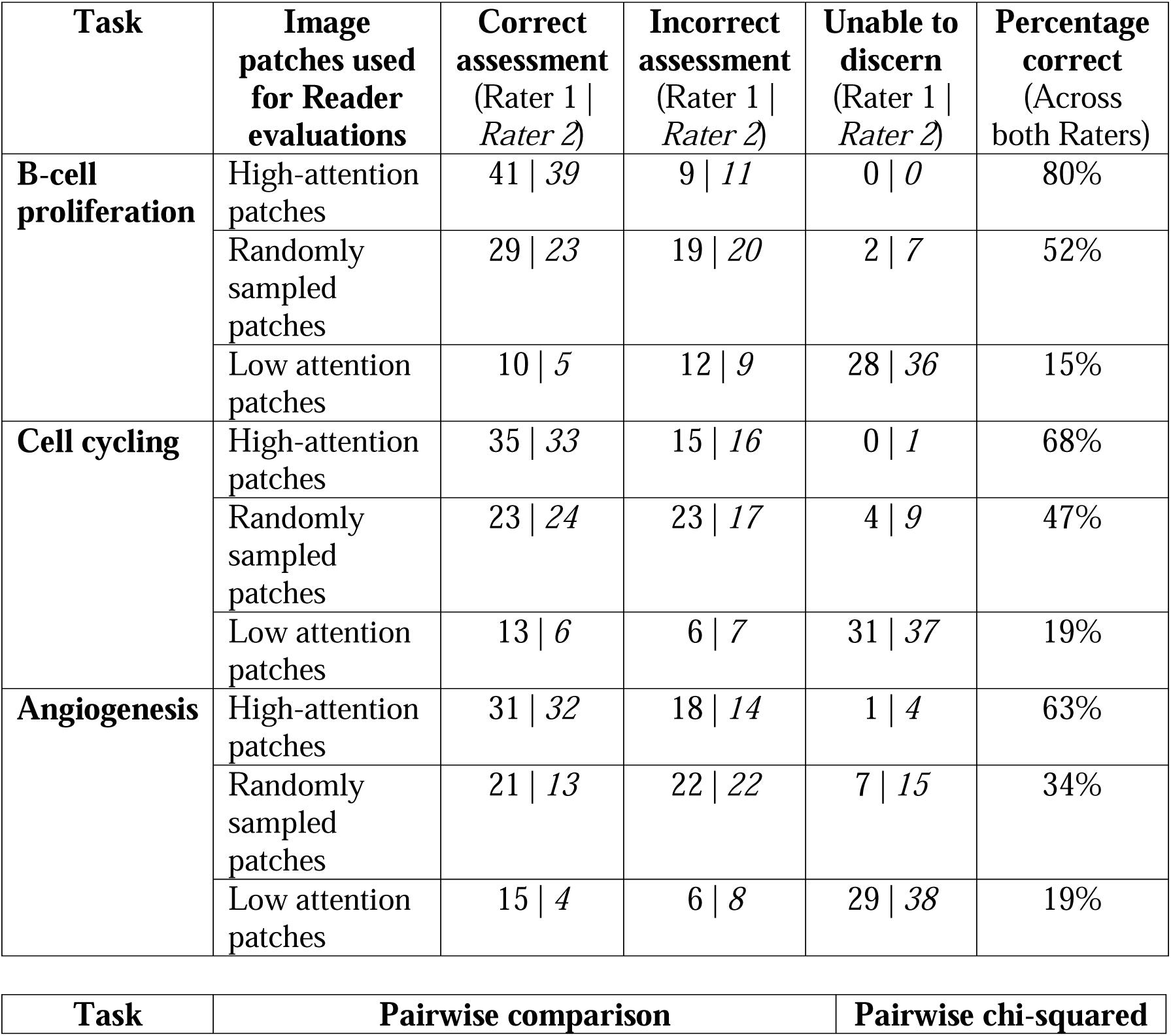
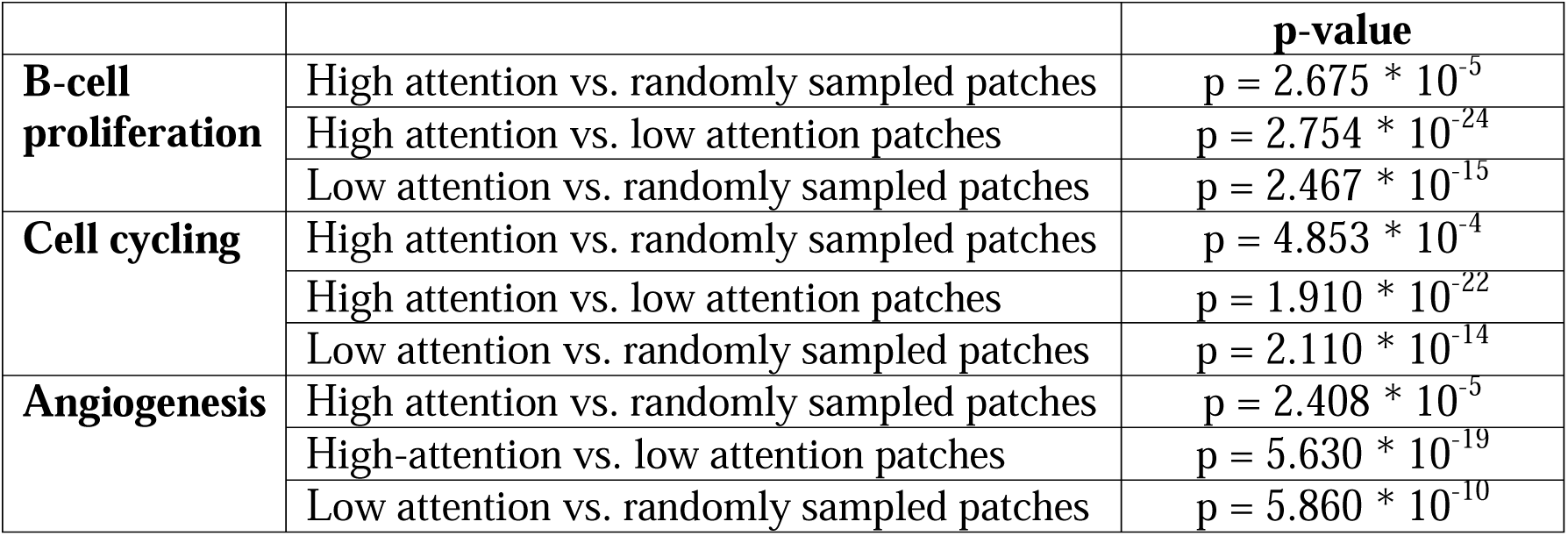
Blinded reader evaluations from two independent raters demonstrate that attention mechanisms identify patches within an H&E WSI enriched for task-specific biology. A) Reader assessments with high-attention patches consistently outperform assessments with randomly sampled or low-attention patches for tasks with visually explicit histologic patterns. B) Pairwise chi-squared comparisons across patch types show significant improvements in reader performance with high-attention patches, and significant reduction in performance with low-attention patches. Together, these results illustrate that attention mechanisms prioritize regions enriched for task-specific histology and de-emphasize less relevant tissue.

| Task | Image patches used for Reader evaluations | Correct assessment (Rater 1 \| Rater 2) | Incorrect assessment (Rater 1 \| Rater 2) | Unable to discern (Rater 1 \| Rater 2) | Percentage correct (Across both Raters) |
| --- | --- | --- | --- | --- | --- |
| B-cell proliferation | High-attention patches | 41 \| 39 | 9 \| 11 | 0 \| 0 | 80% |
|  | Randomly sampled patches | 29 \| 23 | 19 \| 20 | 2 \| 7 | 52% |
|  | Low attention patches | 10 \| 5 | 12 \| 9 | 28 \| 36 | 15% |
| Cell cycling | High-attention patches | 35 \| 33 | 15 \| 16 | 0 \| 1 | 68% |
|  | Randomly sampled patches | 23 \| 24 | 23 \| 17 | 4 \| 9 | 47% |
|  | Low attention patches | 13 \| 6 | 6 \| 7 | 31 \| 37 | 19% |
| Angiogenesis | High-attention patches | 31 \| 32 | 18 \| 14 | 1 \| 4 | 63% |
|  | Randomly sampled patches | 21 \| 13 | 22 \| 22 | 7 \| 15 | 34% |
|  | Low attention patches | 15 \| 4 | 6 \| 8 | 29 \| 38 | 19% |

|  |  |  |
| --- | --- | --- |
| Task | Pairwise comparison | Pairwise chi-squared |

|  |  | **p-value** |
| --- | --- | --- |
| **B-cell proliferation** | High attention vs. randomly sampled patches | $p = 2.675 * 10^{-5}$ |
| | High attention vs. low attention patches | $p = 2.754 * 10^{-24}$ |
| | Low attention vs. randomly sampled patches | $p = 2.467 * 10^{-15}$ |
| **Cell cycling** | High attention vs. randomly sampled patches | $p = 4.853 * 10^{-4}$ |
| | High attention vs. low attention patches | $p = 1.910 * 10^{-22}$ |
| | Low attention vs. randomly sampled patches | $p = 2.110 * 10^{-14}$ |
| **Angiogenesis** | High attention vs. randomly sampled patches | $p = 2.408 * 10^{-5}$ |
| | High-attention vs. low attention patches | $p = 5.630 * 10^{-19}$ |
| | Low attention vs. randomly sampled patches | $p = 5.860 * 10^{-10}$ |

**Table 3:** Model-derived quantitative TME phenotypes stratify for therapeutic outcomes across multi-institutional cohorts for therapeutic regimens with limited predictive biomarkers of efficacy. Values represent median model-derived TME phenotype activity scores (95% confidence interval [CI]) estimated from pre-treatment H&E WSIs within each clinical outcome group. P-values were calculated using two-sided Wilcoxon rank-sum tests. A) <u>KEYNOTE-522 (neoadjuvant pembrolizumab + chemotherapy)</u>: Pre-treatment core biopsies from patients achieving a pathologic complete response (pCR) demonstrated significantly higher model-derived immunogenic activity compared with biopsies from patients with residual disease at surgery. B) <u>Adjuvant dose-dense chemotherapy</u>: Pre-treatment H&E WSIs from patients achieving long-term remission after dose-dense chemotherapy exhibited a trend towards higher immunogenic model-derived phenotypes compared with those from patients who subsequently relapsed.

| Tasks | Model-derived quantitative expression for <u>pathologic complete response cohort</u> (n = 57 patients): Median (95% CI) | Model-derived quantitative expression for <u>residual disease cohort</u> (n = 52 patients): Median (95% CI) | Statistical Significance |
| --- | --- | --- | --- |
| *B-cell proliferation* | 0.831 (0.534, 1.127) | 0.251 (-0.021, 0.523) | p = 0.017 |
| *T-cell mediated cytotoxicity* | 0.365 (0.040, 0.689) | -0.160 (-0.461, 0.141) | p = 0.020 |
| *FoxP3-mediated immunosuppression* | -0.135 (-0.421, 0.150) | 0.454 (0.107, 0.802) | p = 0.004 |
| *Antigen processing and presentation* | 0.543 (0.224, 0.862) | 0.061 (-0.255, 0.376) | p = 0.045 |

B)
| Tasks | Model-derived quantitative expression for <u>Remission cohort</u> (n = 50): Median (95% CI) | Model-derived quantitative expression for <u>Recurrence cohort</u> (n = 29): Median (95% CI) | Statistical Significance |
| --- | --- | --- | --- |
| *B-cell proliferation* | 0.559 (0.414, 0.703) | 0.431 (0.098, 0.642) | p = 0.186 |
| *T-cell mediated cytotoxicity* | 1.763 (1.545, 1.980) | 1.597 (1.170, 2.024) | p = 0.355 |
| *FoxP3-mediated immunosuppression* | 0.585 (0.456, 0.713) | 0.611 (0.179, 0.692) | p = 0.366 |
| *Antigen processing and presentation* | 1.390 (1.155, 1.627) | 1.089 (0.460, 1.276) | p = 0.360 |
| *Cell cycling* | -0.800 (-0.667, -0.932) | -0.880 (-0.761, -0.964) | p = 0.106 |

## DISCUSSION

Mechanistic insights in modern oncology are largely derived from multi-omics assays that are inherently confined to small research cohorts and impractical to deploy in clinical practice. To characterize the breadth of resistance biology across real-world populations, we applied weakly-supervised DL to routine histology to infer *patient-specific* phenotypes and identify the focal tissue regions enriched for this biology. Because these models learn therapeutically relevant states from slide-level labels alone, our approach offers a scalable avenue for extracting biological insights for real world patient populations. Importantly, our work illustrates a potential use of DL-based localization beyond the interpretation of predictions. Whereas attention has typically served as a post hoc check for visually explicit tasks (e.g., confirming that a model attends to diagnostically relevant regions), our findings support its use for prioritizing tissue regions for biological investigation. By identifying regions associated with resistance-related phenotypes, this framework could aid in the discovery of new cellular and molecular mediators of resistance.

Central to this framework is our use of systems-level TME biology, rather than isolated molecular snapshots, for model supervision. Models trained to predict individual gene expression^13,14,27–29^ may provide a less robust representation of functional biology because of functional redundancy (e.g., if one gene is downregulated, others in that pathway may compensate) and the susceptibility of single-gene measurements to technical variance across sequencing platforms^30,31^. Systems-level phenotypes integrate multiple molecular components of a biological program and may therefore provide more stable and functionally informative supervision. Consistent with this rationale, our pathway-level models compared favorably with previously reported single-gene benchmarks^13,14,28,29^ across nearly all TME phenotypes. Moreover, supervision using biological states associated with resistance, rather than outcomes specific to a single treatment, may allow the inferred phenotypes to be applied across therapeutic contexts.

A critical first step was to establish that model-derived phenotypic activity accurately reflected the intended biology. Tissue-matched mIF demonstrated concordance between inferred phenotypes and corresponding mIF-derived cell fractions. While these associations were moderate, perfect agreement was not expected, as these two assays capture related, but not identical, measures: transcriptomic activity reflects a dynamic biological state, whereas mIF provides a direct measure of cellular composition and protein expression. For example, the association between B-cell proliferation and mIF-derived B-cell fractions (PCC: 0.331) is consistent with the model capturing B-cell *proliferative activity* rather than B cell abundance alone. Examination of cases with discordant FoxP3 activity and mIF-derived Treg fractions identified FoxP3 signal in non-Treg compartments (e.g., myeloid or tumor cells). This finding raises the possibility that the model-inferred phenotype reflects a broader FoxP3-associated immunosuppressive state rather than Treg abundance alone. These discrepant cases illustrate how model-derived phenotypes may reveal biological variation that is not captured by cell fractions alone. Collectively, phenotype-consistent associations with orthogonal mIF measurements support the conclusion that the models quantify biologically meaningful TME states.

As a second, orthogonal axis of validation, we asked whether inferred phenotypes had therapeutic relevance. We reasoned that models capturing functional resistance biology should recover established relationships between these phenotypes and therapeutic outcomes. Across multi-institutional cohorts of TNBC patients, model-derived phenotypes were linked to therapeutic outcomes in directions consistent with established determinants of response^20–24^.

Immunogenic signatures were significantly associated with pCR in the KEYNOTE-522 cohort. The more modest associations between immune phenotypes and ddACT outcomes were consistent with the multifactorial and tumor-intrinsic mechanisms underlying chemotherapy response^32,33^ (e.g., DNA repair deficiency, apoptotic activity). Notably, these associations were recovered from core biopsy H&E WSIs, supporting the feasibility of quantifying systems-level biology in tissue-scarce settings typical of clinical oncology. Together with the mIF analyses, these results provide further evidence that these models infer therapeutically relevant biology and support the premise that model attention prioritizes focal regions containing the corresponding biology.

We next tested this premise directly by evaluating whether pathologists could identify phenotype-specific morphology within model-prioritized regions. Attention-based models are highly parameterized and can inadvertently achieve high performance on limited training data by attending to confounders, rather than phenotype-specific biology. In our blinded reader study, pathologist assessments were most accurate using high-attention patches. Conversely, low-attention regions frequently harbored uninformative morphology. These findings provide direct evidence that model-prioritized regions were enriched for recognizable phenotype-specific biology. As attention identifies regions that contribute to phenotype inference rather than defining their underlying biology, these findings position attention-defined regions as a principled starting point for mechanistic investigation. Spatial profiling and functional experiments will be needed to characterize the cellular populations within these regions, nominate candidate mediators of resistance, and establish their functional roles.

The correspondence between weakly supervised learning and the spatial biology of therapeutic resistance may help explain the utility of this approach. The computational challenge of extracting task-relevant signals from thousands of heterogeneous image patches mirrors the biological challenge of resolving resistance-associated signals within tumor tissue. Just as resistance biology may be mediated by localized cellular niches^1–5^, the histologic signals associated with therapeutic resistance may be confined to a subset of image patches. Bulk profiling assays, by contrast, average molecular signals across heterogeneous tissue and may obscure focal biological programs that contribute to patient-level phenotypes. Phenotype-directed localization therefore links patient-level states of therapeutic efficacy to the focal tissue regions that contribute most strongly to model inference.

Our framework is complementary to cross-modal foundation models that align histopathology with ST or multiplexed imaging^34–37^. Models such as GigaTIME^37^ and STORM^36^ can derive high-resolution spatial readouts from routine H&E WSIs but require paired histologic and spatial molecular data for training. Our weakly supervised approach addresses a different objective. Rather than reconstructing predefined molecular measurements throughout the tissue, it uses patient-level labels to prioritize the focal regions most strongly associated with a biological or therapeutic state. Because patient-level labels are more widely available than paired spatial data, this approach could be applied across large histopathology cohorts. The breadth of resistance biology captured by our models further motivates <u>resistance-directed</u> <u>localization</u>, in which therapeutic outcomes could serve as supervision to identify focal regions associated with response or resistance. These approaches may operate sequentially: resistance-directed localization would first identify the tissue regions where focused investigation may be most informative, after which spatial profiling resolves the cellular populations and molecular programs within those regions.

Several limitations warrant consideration. First, our models used AM-SB, an ABMIL-based architecture that treats WSI patches as independent instances and therefore does not explicitly capture spatial relationships across adjacent tissue regions^38^. Although transformer-based models were evaluated, they did not outperform AM-SB in the present cohort, potentially because learning more complex spatial interactions requires larger training datasets^39,40^. The optimal architecture for localizing therapeutically relevant biology therefore remains uncertain. Future studies should compare alternative localization strategies, including multi-head attention pooling, spatially informed architectures, and concept-based approaches (e.g., ProtoMIL^41^).

Second, orthogonal validation was not available for every modeled phenotype. Metabolic programs were not evaluated using mIF, therapeutic outcomes, or blinded pathologist assessments. Unlike immune phenotypes, these programs lack readily available protein-level markers or recognizable histologic correlates, and their relationships with therapeutic efficacy are less firmly established. Additional molecular and spatial validation will be needed to determine whether this framework extends to metabolic and other less visually apparent biological programs. Third, our analyses were limited to breast cancer, and the orthogonal validation cohorts were relatively modest in size (e.g., 25 patients for mIF; 188 patients across therapeutic outcomes cohorts). Validation in larger cohorts and additional cancer types will therefore be needed. Finally, this study does not establish that molecular interrogation of model-prioritized regions will uncover causal mediators of resistance. Nonetheless, the present results motivate targeted spatial profiling to characterize the cellular populations and molecular programs within model-prioritized regions, followed by functional experiments to establish the roles of nominated candidates.

In summary, we demonstrate that weakly supervised DL can infer therapeutically relevant TME phenotypes from routine histology and localize the focal tissue regions enriched for the corresponding biology. Given emerging evidence that resistance is often mediated by focal niches within the TME, this approach provides a scalable strategy for focusing multi-omic investigation on regions linked to resistance-associated biology. In future work, therapeutic outcomes could serve as supervision to identify tissue regions associated with response or resistance. Spatial profiling of these model-prioritized regions could then nominate cellular and molecular mediators for functional validation across large clinical populations. Herein, DL can bridge large clinical data repositories with spatially resolved discovery of resistance biology.

## METHODS

### Datasets

The training dataset was comprised of H&E WSIs, patient-matched bulk RNA-sequencing data, and associated clinical data obtained from The Cancer Genome Atlas – Breast Cancer and cBioPortal for Cancer Genomics^42,43^. In total, the dataset included 3111 WSIs from 1098 breast cancer patients. All WSIs were scanned at 40x magnification using Aperio scanners. All data were generated as part of routine clinical care and not within the context of a prospective clinical trial.

The ground truth was derived from the bulk RNA-sequencing data for each WSI in our training set. Normalized gene expression values for the TCGA cohort were obtained from cBioPortal. TME phenotypes were defined using pre-specified gene sets derived from gene ontology-based pathway analyses^44^. We selected ten TME phenotypes, spanning tumor-intrinsic, immune, and metabolic programs, each of which has been implicated in therapeutic resistance across diverse tumor types and treatment modalities^45–47^. For each patient-TME phenotype pair, we computed gene set enrichment scores using single-sample gene set enrichment analysis (ssGSEA). ssGSEA converts an individual sample’s gene expression profile into a gene set-level enrichment profile, yielding a quantitative score that reflects the coordinated up-regulation (positive values) or down-regulation (negative values) of genes within a given biological pathway^48,49^.

To evaluate model generalizability and clinical relevance, we curated two further cohorts of H&E slides from breast cancer patients treated at three institutions: Massachusetts General Hospital (MGH), Brigham & Women’s Hospital (BWH), and the Ohio State University (OSU). All slides were scanned at 40x magnification using routine clinical workflows on either Lecia Aperio GT450 DX (MGH, BWH) or Philips Intelisite Ultrafast scanners (OSU). The first cohort (n = 121 patients from MGH, BWH, OSU) consisted of H&E WSIs of pre-treatment core biopsies from triple-negative breast cancer patients who went on to receive the KEYNOTE-522 regimen^19^. This regimen consists of neoadjuvant pembrolizumab, paclitaxel, and carboplatin, followed by definitive surgery. Clinical outcomes were binarized based on surgical pathology following neoadjuvant therapy. Patients were classified as achieving a pathologic complete response (pCR) if no residual tumor was identified in the breast or sampled lymph nodes, and as having residual disease if any residual invasive tumor was present at definitive surgery. A second cohort (n = 78 patients from MGH and BWH) consisted of pre-treatment H&E WSIs from patients with triple-negative breast cancer who subsequently received adjuvant anthracycline- and taxane-based chemotherapy^50^. Clinical outcomes were binarized based on longitudinal follow-up: patients were classified as remission if they remained disease-free for at least seven years of follow-up, and as recurrence if they developed loco-regional or distant disease recurrence. For both cohorts, there was no associated tabular clinical or demographic data. All WSIs were scanned at 40x magnification using routine clinical workflows. Automated quality control checks for artifact burden and tissue sufficiency were applied, followed by pathology review to confirm tumor adequacy. These WSIs were processed with the same pre-processing pipelines used for model development.

### WSI Processing

#### Segmentation, quality control, and patching

WSI pre-processing followed a protocol consistent with prior work in WSI DL-based classification^6,8,14,35,51,52^. Each WSI underwent quality assessment using the HistoQC framework, an open-source tool for automated WSI quality control that evaluates image quality metrics, artifact detection, and tissue segmentation^53^. HistoQC was applied using the default parameters, yielding curated segmentation masks distinguishing tissue from background. Using this mask, each WSI was partitioned into non-overlapping 256 x 256-pixel patches at 40x magnification.

#### Label-preserving data augmentation strategies

To augment the training data, we implemented label-preserving strategies designed to improve model generalization by simulating common sources of variability in histopathology.

1. **HED Color Augmentation**: To address the color and intensity variability arising from differences in tissue preparation and staining within batches of WSIs from different sites, we applied stain-aware color perturbations at the tile level. Image patches were first converted from RGB to the Hematoxylin-Eosin-DAB (HED) color space, which separates the contributions of individual stains and enables targeted manipulation. The intensities of the hematoxylin-and-eosin channels were independently scaled by a factor sampled uniformly from 0.8 to 1.2, a range consistent with established approaches for simulating realistic staining variability without introducing visually implausible artifacts that could negatively impact model training^54^. This process was used to pre-compute four augmented versions of each slide.
2. **Tile Shifting Augmentation**: To enhance spatial invariance and robustness to minor variations introduced during slide digitization, we implemented a tile-shifting strategy that alters the origin of the tiling grid applied to each WSI. Specifically, we generated three additional sets of tiles by shifting the grid by half a patch’s length (128 pixels) horizontally, vertically, and along both diagonal directions. This strategy produces alternate patch configurations with overlapping but distinct spatial contexts, encouraging the model to learn more generalizable morphological features.
3. **Geometric transforms**. We applied standard geometric transformations at the tile level, including horizontal and vertical flips, to further increase morphological diversity.

During the training loop, we employed a two-stage probabilistic sampling scheme to introduce feature-level variability on-the-fly, balancing data diversity with the computational constraints of multiple-instance learning workflows that rely on pre-computed features. At the slide level, each WSI had a 20% probability of being selected for augmentation. For selected slides, a second sampling step was performed at the tile level, in which each tile had a 50% probability of having its feature vector replaced. For tiles selected for replacement, one of the corresponding pre-computed augmented feature vectors was chosen uniformly at random. This stochastic augmentation strategy ensured continuous exposure to diverse and iterative augmented features during training without requiring repeated feature extraction.

#### Feature extraction

We extracted feature vectors from each patch using three different feature extractors. First, we used CONtrastive learning from Captions for Histopathology (CONCH)^52^, a vision-language foundation model for histopathology pretrained on paired histopathology image and captions. Second, we evaluated Pathology Language–Image Pretraining (PLIP), a multimodal foundation model jointly trained on image-text pairs from the OpenPath dataset^55^. Third, we used ResNet50^56^, a convolutional neural network (CNN) pre-trained on ImageNet.

To identify the best feature representation for downstream analysis, we trained a binary classification model using each set of feature vectors and compared performance across the three feature extractors. Based on these comparisons, the best-performing feature extractor was used for all subsequent experiments.

### H&E WSI model training

Our model was trained to quantify TME phenotypes using two approaches: binary classification and regression. This dual modeling strategy was motivated by clinical considerations: while binary classification enables coarse stratification of phenotypic activity (e.g., up- or down-regulation), continuous estimates of TME phenotype expression may better support precision-based decision making. We did this because in clinical practice, binary classification is too coarse a classification by which to make precision-based decisions; hence it is helpful to know the exact amount of over- or under-expression. For binary classification, ssGSEA scores were binarized such that negative values (pathway down-regulation) were assigned label 0 and positive values (pathway up-regulation) were assigned the label 1. Therefore, the output for binary classification models was either label 0 or 1. For regression, our model directly predicted normalized ssGSEA scores as continuous outcomes. A random data split, at the patient level, allocated 70% to training, 15% to validation, and 15% to test sets, ensuring that all data from a given patient were confined to the same split. Data within each partition was balanced for ER, PR, and HER2 receptor profiles.

For the models using only WSIs as the sole input modality, we employed the attention-based multiple-instance learning architecture (AM-SB) described in our prior work for phenotyping^17^. AM-SB is a modification of the single-branch CLAM framework (CLAM-SB)^8^. Unlike the original CLAM-SB architecture, AM-SB removes instance-level clustering under a mutual exclusivity assumption, which is inappropriate for TME phenotypes that may co-occur spatially within the same slide. In addition to AM-SB, we also evaluated a vision transformer (ViT)-based MIL architecture (TransMIL) to assess whether modeling long-range spatial dependencies between WSI patches improved performance for TME phenotyping^57^.

Patch-level feature vectors were extracted using the CONCH encoder^52^, the best-performing feature extractor, were provided as input to the model. These features were first projected from their original dimensionality into a 512-dimensional latent space using a fully connected layer with a ReLU activation function. These transformed features were then processed by a gated attention mechanism that assigns an attention weight to each patch, reflecting its contribution to the slide-level representation. Attention-weighted aggregation across the slide yielded a single 512-dimensional slide-level feature vector, which was passed to a task-specific prediction head. For TransMIL models, patch embeddings were processed through transformer layers to capture contextual relationships across spatially separate instances, followed by slide level aggregation and a task-specific prediction head.

For binary classification, the prediction head consisted of a fully connected linear layer that performs a linear mapping from the 512-dimensional slide-level feature vector to a 2-dimensional logit space, followed by a softmax activation. Models were trained using a slide-level cross-entropy loss function. To mitigate class imbalance at the slide level, we employed a class-weighted sampling strategy.

For regression tasks, the classification head was replaced with a single linear neuron mapping the slide-level feature vector to a scalar output, with no final activation function applied. Because regression targets are continuous and defined at the slide level, instance-level clustering supervision was not applied. Regression models were trained using slide-level mean squared error loss.

All models were trained for 300 epochs using the adaptive moment (Adam) optimizer with a learning rate of 2 x 10^-4^ and a weight decay of 1 x 10^-5^. To prevent overfitting, we applied a dropout rate of 0.25, applied to the units in the network layers, and used early stopping with a patience of 20 epochs, monitored on the validation loss. Early stopping was not initiated before a minimum of 25 training epochs had been completed. Model selection was performed based on validation performance, and the best-performing checkpoint was used for evaluation on the held-out test set.

### Training of classification models using clinico-genomic data

We trained binary classification models using clinico-genomic metadata routinely used in clinical trial stratification as model inputs. The clinical variables evaluated in this study included: age, sex, race, American Joint Committee on Cancer (AJCC) staging, histological subtype, PAM50 molecular subtype, estrogen receptor (ER) status, progesterone receptor (PR) status, and human epidermal growth factor 2 (HER2) status. Age was discretized into categorical bins: 18-30; 30-40; 40-50; 50-60; 60+. All variables (both originally categorical and discretized) were encoded using one-hot representations prior to model input.

To establish a baseline for the ability of standard-of-care clinical data to measure therapeutically relevant biology, we trained binary classification models, using only tabular clinico-genomic data. These experiments were conducted prior to multimodal model development to quantify the independent predictive contribution of these tabular clinical variables. We evaluated three supervised learning approaches for clinico-genomic-only modeling: random forest, support vector machine (using a radial basis function non-linear kernel), and tabPFN, a transformer-based foundation model for tabular data pretrained on synthetic classification tasks^58^.

### Multimodal model training strategies

As the optimal strategy for combining H&E WSI with clinico-genomic data remains an active area of investigation, we evaluated two fusion approaches: cross-attention based intermediate fusion and late fusion of modality-specific predictions.

Our first approach was an intermediate fusion strategy adapted from the SurvPath architecture^18^. In this framework, we modified the original transcriptomics-based biological pathway tokenization to accommodate tabular clinico-genomic features, while retaining the cross-attention mechanism to integrate clinical and whole-slide image (WSI) representations. The cross-attention layer enables the model to learn inter-modality interactions that may be relevant for the downstream task. To address the dimensionality mismatch between CONCH-derived WSI features (512 dimensions) and the lower-dimensional clinical feature vectors (typically 1-10 dimensions after encoding), we applied a learnable multilayer perceptron (MLP) to project clinical features into a higher-dimensional embedding space (expanded by a factor of 100), ensuring compatibility with the WSI feature representations prior to fusion. Models were trained for 300 epochs using mean-squared error loss and the Adam optimizer, with a learning rate of 2 × 10^-4^ and weight decay of 1 × 10^-5^. The model checkpoint achieving the lowest validation loss was selected for final evaluation.

Our second approach was a late fusion strategy^51^, in which modality-specific models were trained independently and their outputs combined at the prediction level. Predictions from the WSI-only and clinico-genomic-only models were aggregated using an MLP consisting of an input layer with two neurons; two hidden layers with four and two neurons, respectively; and a single-neuron output layer. This fusion network was trained for 300 epochs using mean-squared error loss and the Adam optimizer, with identical learning rate and weight decay settings as above. Model selection was again based on validation loss.

### Evaluation of model performance

For binary classification on H&E WSI-only and multimodal models, performance was evaluated using the area under the receiver operating characteristic curve (AUROC) metric. For regression models predicting continuous ssGSEA-derived phenotypes, we report Pearson correlation coefficient (PCC) between predicted and ground-truth scores. The clinico-genomic-only models were evaluated using AUROC. To quantify uncertainty, we performed bootstrap resampling with 1000 iterations. For each bootstrap run, performance metrics were recomputed on the resampled test sets, and a median and 95% confidence interval was reported across bootstrap distributions.

### Ablation studies

To quantify the impact of different categories of clinico-genomic information within the clinico-genomic and multimodal model settings, we conducted ablation studies by training models using predefined subsets of clinical features. Three clinical feature sets were considered: (i) **All** (age, gender, race, AJCC stage, histologic subtype, PAM50 subtype, ER status, PR status, and HER2 status); **Demographics** (age, gender, race); and **Molecular** (PAM50 subtype, ER status, PR status, and HER2 status).

### Computational environment

Model training was conducted on a SLURM-managed high-performance computing (HPC) cluster, using single NVIDIA RTX 8000 GPUs with 4 CPU cores and 200 GB of RAM. The data loading pipeline utilized 4 parallel workers to ensure efficient data throughput. For neural network training and optimization, we used the PyTorch framework and for WSI manipulation, we used OpenSlide. The code related to the implementation of this paper is publicly available in a GitHub repository: https://github.com/QTIM-Lab/tme-wsi.

### Visualization of high- and low-attention regions on H&E WSIs

Attention weights were produced by the attention-based MIL framework during slide-level inference^8^. For each WSI and TME phenotype, attention scores were assigned to individual image patches and normalized within each WSI. These scores reflect the relative contribution of each patch to the model’s final prediction.

Next, to interpret model predictions and assess spatial correlates of TME phenotypes within WSIs, attention maps were generated. These maps were visualized by projecting patch-level attention scores back onto their spatial coordinates within the original WSIs. Heatmaps were generated using a continuous color scale, with regions in red corresponding to high attention weights and regions in blue corresponding to low attention weights. As distinct TME phenotypes correspond to different morphologic patterns or cell populations, attention maps were generated separately for each phenotype to enable phenotype-specific interpretation. High-attention regions were qualitatively reviewed by a board-certified pathologist to identify recurring spatial or morphologic patterns associated with specific phenotypes.

### Model validation on therapeutic contexts with multi-institutional external data

To evaluate the generalizability and clinical relevance of our model, we curated multi-institutional cohorts of pre-treatment H&E WSIs from breast cancer patients treated at Massachusetts General Hospital (MGH), Brigham & Women’s Hospital (BWH), and the Ohio State University. None of these WSIs overlapped with the training or internal validation datasets. For our KEYNOTE-522 cohort, inference was performed on pre-treatment H&E core biopsy WSIs, using the H&E WSI-only regression model. These model-derived phenotypes were compared between pCR and residual disease (thus implying ICI resistance) outcome groups. For the ddACT cohort, inference was performed using the H&E WSI-only regression model. These model-derived phenotypes were compared between remission and relapse outcome groups. For both clinical use cases, differences in model-derived phenotype scores across the outcome groups were assessed using two-sided Mann Whitney U tests. Clinical outcome information was not used during model training or inference.

### Reader study to assess biological fidelity of attention maps

To evaluate whether attention-based learning reliably captured task-specific biology, we conducted a blinded reader study by two board-certified pathologists. Fifty H&E WSIs were randomly selected from the held-aside test set. Using our H&E WSI classification model, inference was performed for three TME phenotypes selected *a priori* based on their known association with histological patterns identifiable on routine H&E review. These phenotypes included: B-cell proliferation, epithelial-mesenchymal transition, and angiogenesis. For each TME phenotype, attention scores were extracted for all image patches within each WSI. From each slide and task, three patch groups were algorithmically selected: the five patches with the highest attention scores, the five patches with the lowest attention scores, and five randomly sampled patches. Patch selection was fully automated and performed without human input.

These patch sets were presented to both blinded pathologists in randomized order, blinded to attention scores and model predictions. For each set of five patches, the pathologist was asked to determine whether the observed histologic features were most consistent with high expression, low expression, or not assessable for the queried phenotype (e.g., insufficient tissue, adipose tissue, background artifact). Because high-attention patches are expected to most strongly reflect task-specific biology, we hypothesized that pathologist assessment of TME phenotypes would be most accurate using the high-attention patches, compared to randomly sampled or low attention patches. Conversely, as low attention patches are expected to have minimal biological relevance to the classification task, we anticipated a higher frequency of “not assessable” determinations within this group.

### Multiplexed immunofluorescence analysis

#### Staining and Imaging

Highly multiplexed spatial proteomics was performed on formalin-fixed, paraffin-embedded (FFPE) Triple-Negative Breast Cancer (TNBC) tissue microarrays using the CODEX/PhenoCycler platform^59^. Tissues were stained with a custom 19-marker panel of oligonucleotide-barcoded antibodies from Akoya Biosciences. These anti-human antibodies targeted CD3e, CD4, CD8, CD11b, CD11c, CD14, CD20, CD21, CD38, CD45, CD56, CD68, CD79a, CD206, FoxP3, Granzyme B (GZMB), HLA-DR, iNOS, and Pan-Cytokeratin (PanCK). Automated cyclic imaging was performed to iteratively hybridize, image, and strip fluorescently labeled reporter oligonucleotides corresponding to the antibody barcodes.

#### Image Processing and Single-Cell Segmentation

Computational processing of the raw multi-channel QPTIFF images was executed using the SPACEc (Structured Spatial Analysis for Codex Exploration) Python framework^60^. Within the SPACEc pipeline, deep learning-based whole-cell segmentation was performed using the integrated Mesmer algorithm, utilizing the DAPI nuclear counterstain and membrane markers to define cell boundaries. Single-cell mean fluorescence intensities for all 19 markers were quantified and extracted into a single-cell expression matrix.

#### Data Preprocessing and Batch Correction

Downstream preprocessing and clustering were also managed within SPACEc and the Scanpy ecosystem. Single-cell marker intensities were Z-score normalized per slide. To correct for batch effects across the four distinct tissue microarray slides, Harmony integration was applied to the principal components^61^.

#### Unsupervised Clustering and Cell-Type Annotation

To prevent the artificial fragmentation of functional cell states, clustering was strictly limited to a 19-marker structural lineage sub-panel. Cells were clustered using the Leiden algorithm via a k-nearest neighbor (k-NN) graph built on the Harmony-corrected embeddings. Clusters were manually annotated into specific immune and tumor cell lineages based on canonical marker expression profiles. Finally, absolute cell counts and relative cellular fractions for these specific populations were extracted per tissue region to serve as the ground-truth cellular metrics for downstream correlation analysis.

## Acknowledgments

The authors acknowledge helpful feedback by Gromit Persimmon Perrino. <u>Funding</u>: William G. Kaelin, Jr., M.D., Physician-Scientist Award of the Damon Runyon Cancer Research Foundation (PST-36-21; A.E.K.)

American Association for Cancer Research Breast Cancer Research Fellowship (A.E.K.)

American Brain Tumor Association Basic Research Fellowship In Honor of Paul Fabbri (A.E.K.)

American Society of Clinical Oncology/Conquer Cancer Young Investigator Award (A.E.K.)

American Cancer Society Institutional Research Grant (IRG-21-130-10; A.E.K.)

American Academy of Neurology Career Development Award (A.E.K.) National Cancer Institute K12CA090354 (A.E.K.)

National Institute of Biomedical Imaging and Bioengineering K08EB037077 (A.E.K.)

Breast Cancer Research Foundation (to D.C.S.)

Victoria’s Secret Global Fund for Women’s Cancers Career Development Award in Partnership with Pelotonia and AACR (F.G.)

Quanta Computer, Inc. (A.V.D.; J.V.G.).

## Author Contributions

**Conceptualization:** A.E.K., C.P.B., T.G.

**Methodology:** T.G., D.P., C.P.B., A.E.K.

**Formal analysis:** T.G., D.P., C.M.P., T.L., J.A.H., V.Z., F.G., C.P.B., A.E.K.

**Investigation:** T.G., A.J.I., K.T.F., J.V.G., A.E.K., C.P.B.

**Visualization:** T.G., A.E.K., C.P.B.

**Writing – original draft:** A.E.K., C.P.B., T.G.

**Writing – review and editing:** All authors.

**Project administration:** A.E.K., C.P.B.

**Funding acquisition:** A.E.K., C.P.B., D.C.S., F.G., A.V.D., J.V.G.

## Competing Interests Statement

SAW reports: consulting and advisory board engagements at Foundation Medicine, Veracyte, Hologic, Eli Lilly, Biovica, Pfizer, Arvinas, Puma Biotechnology, Novartis, AstraZeneca, Genentech, Regor Therapeutics, Stemline/Menarini, Gilead, Celcuity, Precede Biosciences, Halda Therapeutics, Boundless Bio; education and speaking engagements at Eli Lilly, Guardant Health, 2ndMD; and institutional research support (to MGH) from Genentech, Eli Lilly, Pfizer, Arvinas, Nuvation Bio, Regor Therapeutics, Sermonix, Puma Biotechnology, Stemline/Menarini, Phoenix Molecular Designs. KTF is on the board of directors at Strata Oncology, Antares Therapeutics, Gyges Oncology, Khora Therapeutics, Monimoi Therapeutics; the scientific advisory board at Apricity, Tvardi, xCures, ALX Oncology, Karkinos, Soley, Flindr, Alterome, intrECate, PreDICTA, Tasca, Zola, Synthetic Design Labs, and consults for Nextech, Takeda, Transcode Therapeutics.

## Data Sharing Statement

Our model was trained using publicly available data from The Cancer Genome Atlas and the cBioPortal for Cancer Genomics. The raw clinical, H&E WSI, and multiplexed immunofluorescence data in our external validation set are protected due to patient privacy laws. Any requests for raw and analyzed data should be sent in writing to Albert Kim and will be reviewed by the DF/HCC Institutional Review Board (IRB). Pending IRB approval, deidentified data then will be transferred to the inquiring investigator in an expeditious fashion over secure file transfer.

## Code Availability

Our repository for data analysis is located at: https://github.com/QTIM-Lab/tme-wsi.

## List of Supplementary Materials

**Supplemental Figure 1:**
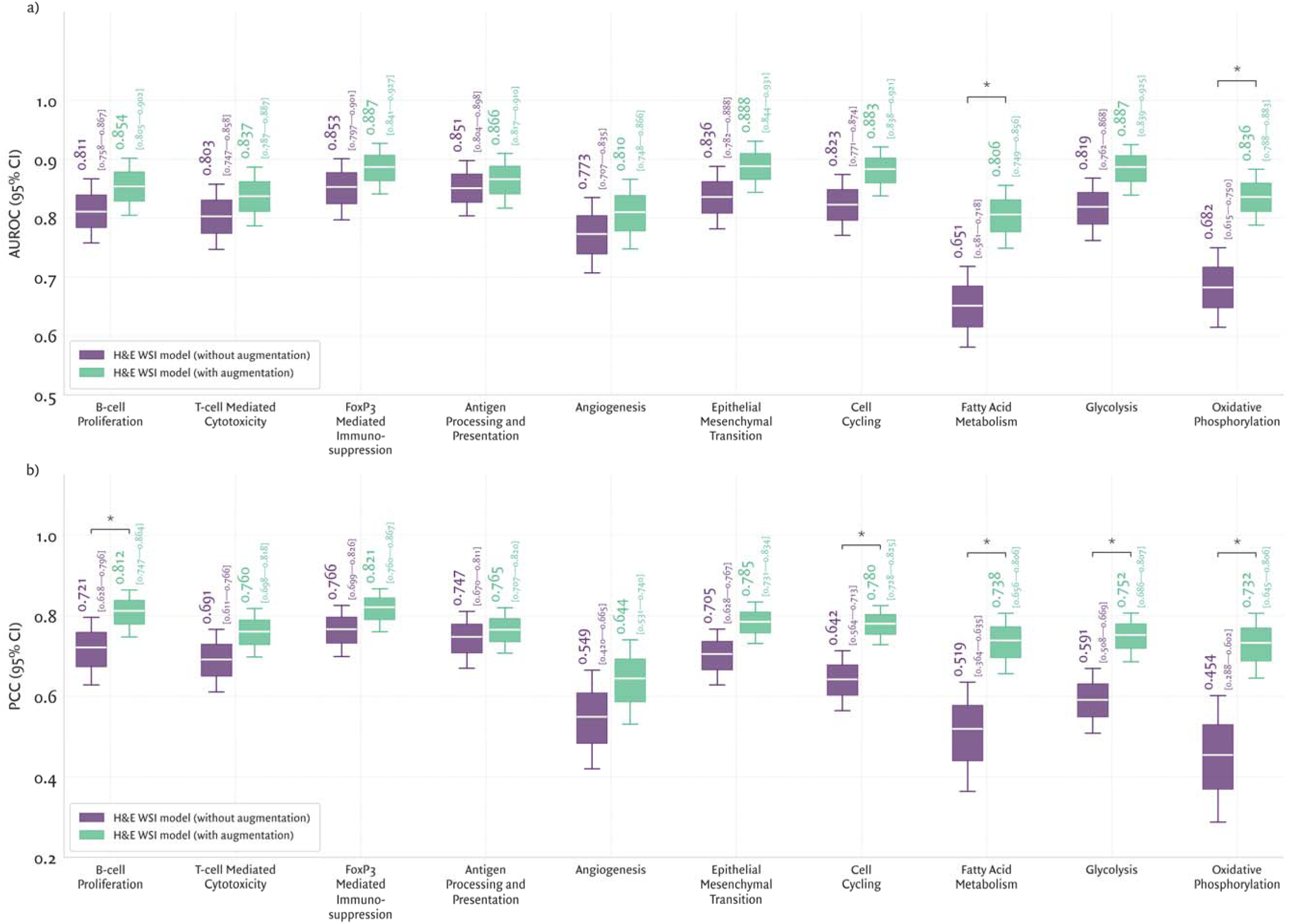
Patch-level image augmentation improves H&E WSI model performance across multiple TME phenotyping tasks. A) AUROC (binary classification) and B) PCC (regression) for ten TME phenotypes, spanning immune, tumor, and metabolic programs. Models compared include H&E WSI without augmentation (purple) and H&E WSI with patch-level augmentation (teal). Box-and-whisker plots display the median and 95% confidence intervals estimated via bootstrap resampling across the held-aside test set. Asterisks include significant differences, relative to the H&E WSI model. Here, patch-level augmentation yields consistent, and often significant, performance gains, suggesting that label-preserving transformations that emulate histologic variability enhance performance of WSI-based TME phenotyping.

**Supplemental Figure 2:**
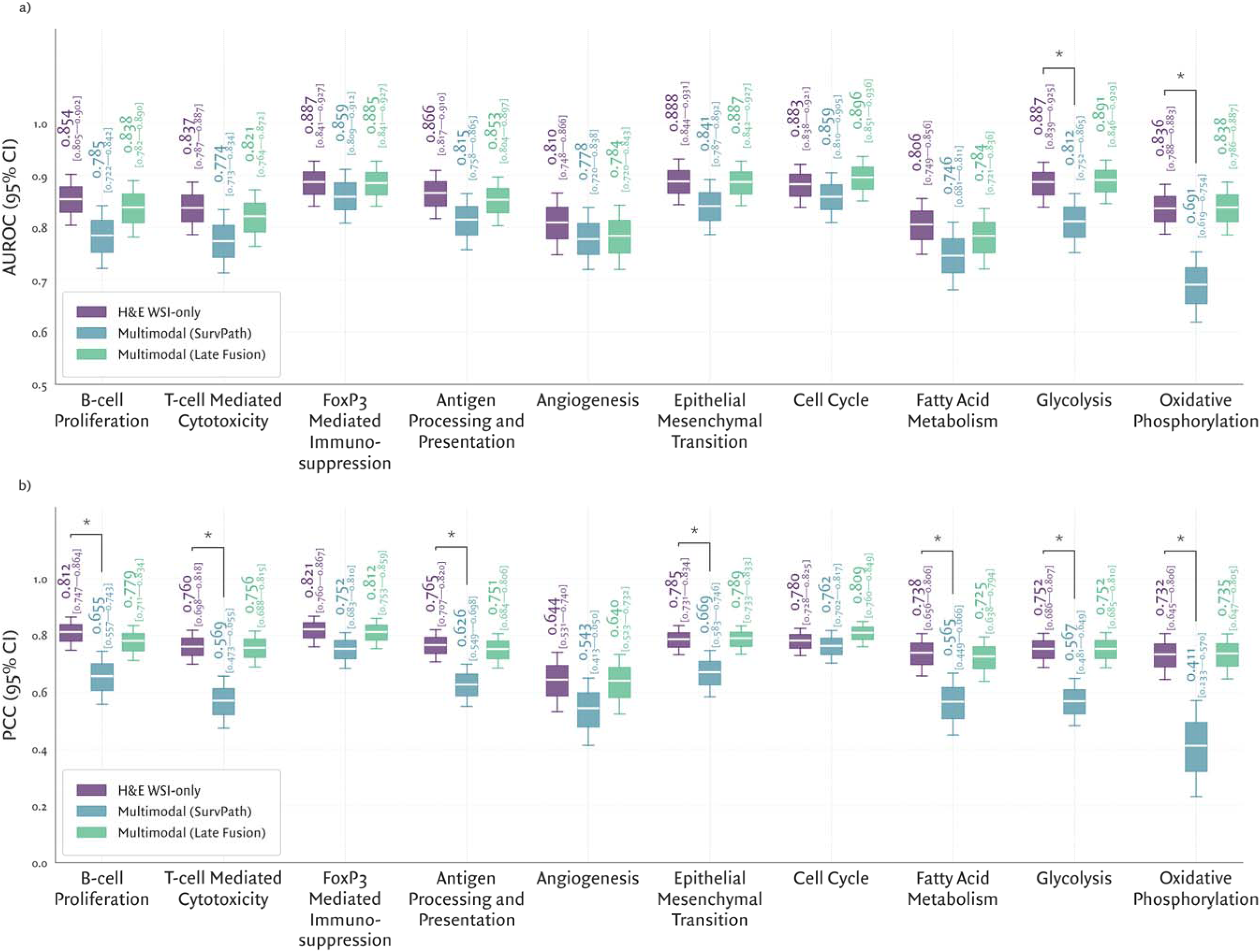
Multimodal integration of H&E WSIs with tabular clinico-genomic data does not improve quantification of systems level TME biology. A) AUROC (binary classification) and B) PCC (regression) for ten TME phenotypes, spanning immune, tumor, and metabolic programs. Models include: H&E WSI (purple), multimodal cross-attention (teal), and multimodal late fusion (green). Box-and-whisker plots display the median and 95% confidence intervals estimated via bootstrap resampling across the held-aside test set. Asterisks include significant differences, relative to the H&E WSI model. Across all tasks, multimodal models do not yield performance gains and frequently underperform the H&E WSI model baseline, suggesting that tabular clinico-genomic data provide limited complementary information for TME phenotyping, or are redundant with signals already encoded within H&E WSIs.

**Supplemental Figure 3:**
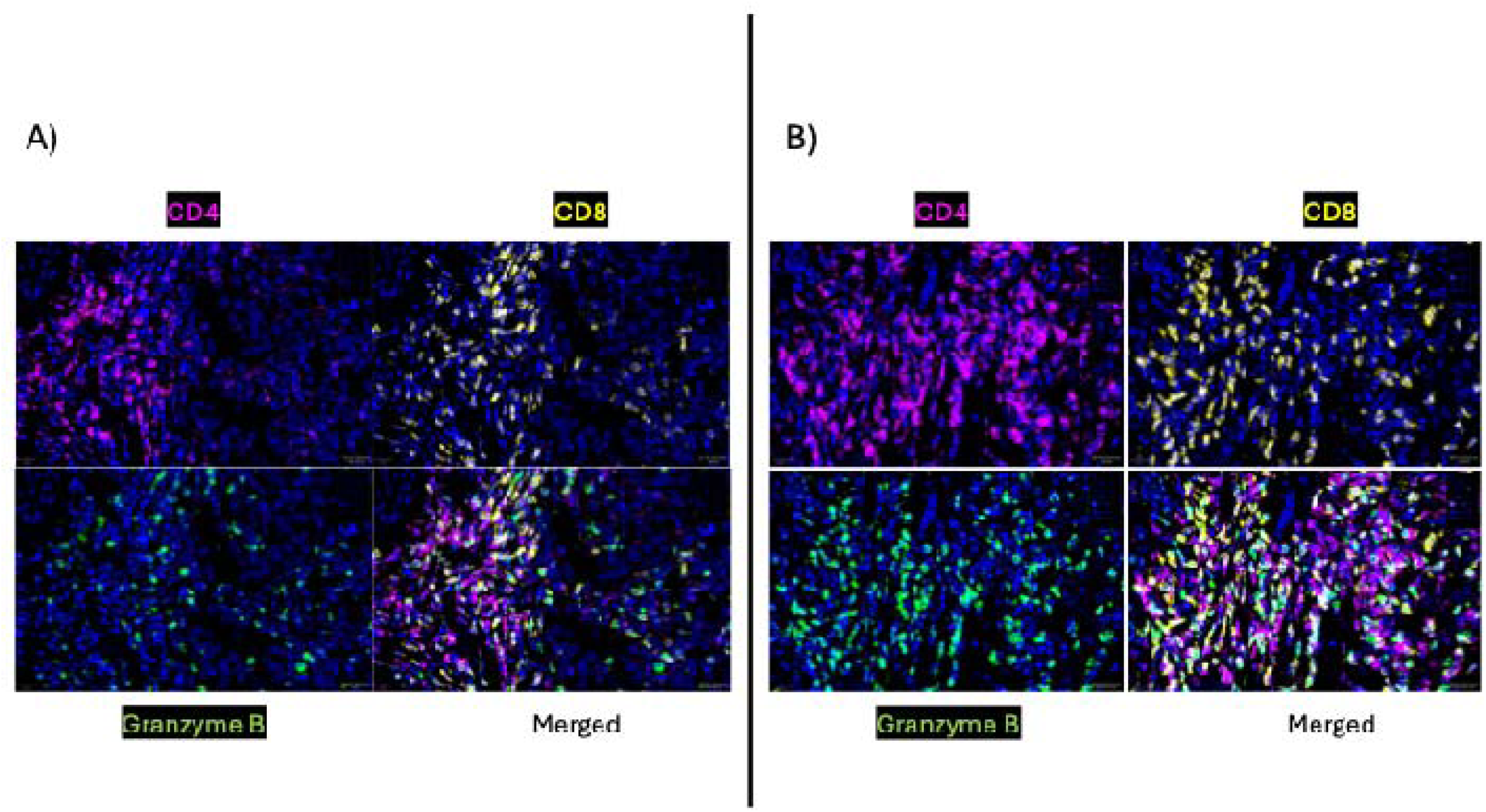
Characterization of cytotoxic T-cells via multiplexed immunofluorescence. (A, B) Exemplar multiplex immunofluorescence images from two distinct patients illustrating the distribution of cytotoxic T-cell infiltrates within the TME. Individual and merged panels display CD4 (magneta), CD8 (yellow), granzyme B (GZMB, green), and DAPI (blue). The merged images demonstrate the presence of cytotoxic T-cell populations, identified by the co-localization of the effector protease GZMB with CD4^+^ or CD8^+^ lymphocytes.

**Supplemental Figure 4:**
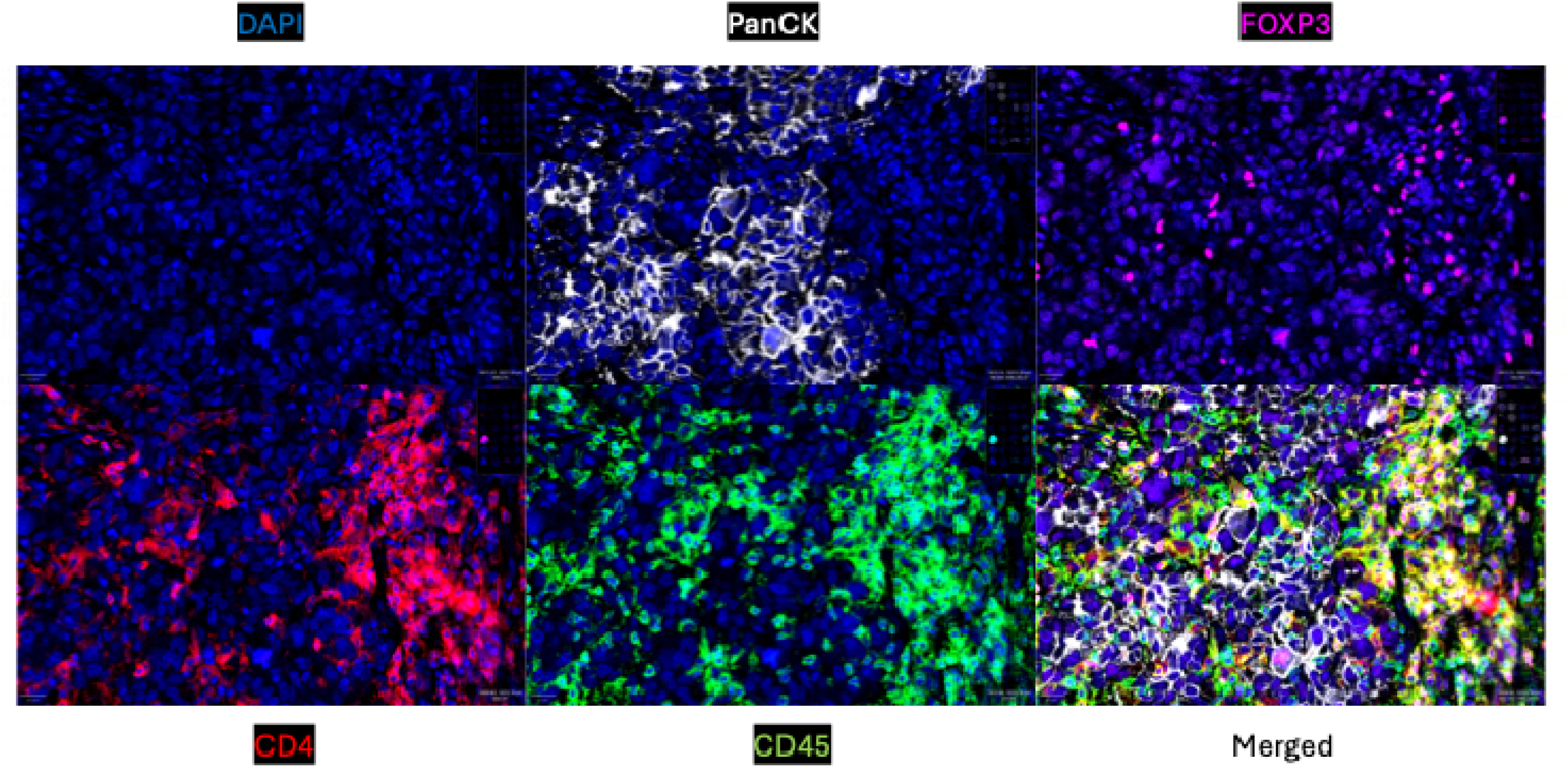
Representative multiplex immunofluorescence images of regulatory T cells (Tregs) within the tumor microenvironment. Sequential panels display individual channels for DAPI (blue, nucleus), PanCK (white, tumor), FoxP3 (magneta), CD4 (red), and CD45 (green). The merged panel demonstrates definitive identification of Tregs by the distinct triple expression of FoxP3, CD4, and CD45.

**Supplemental Figure 5:**
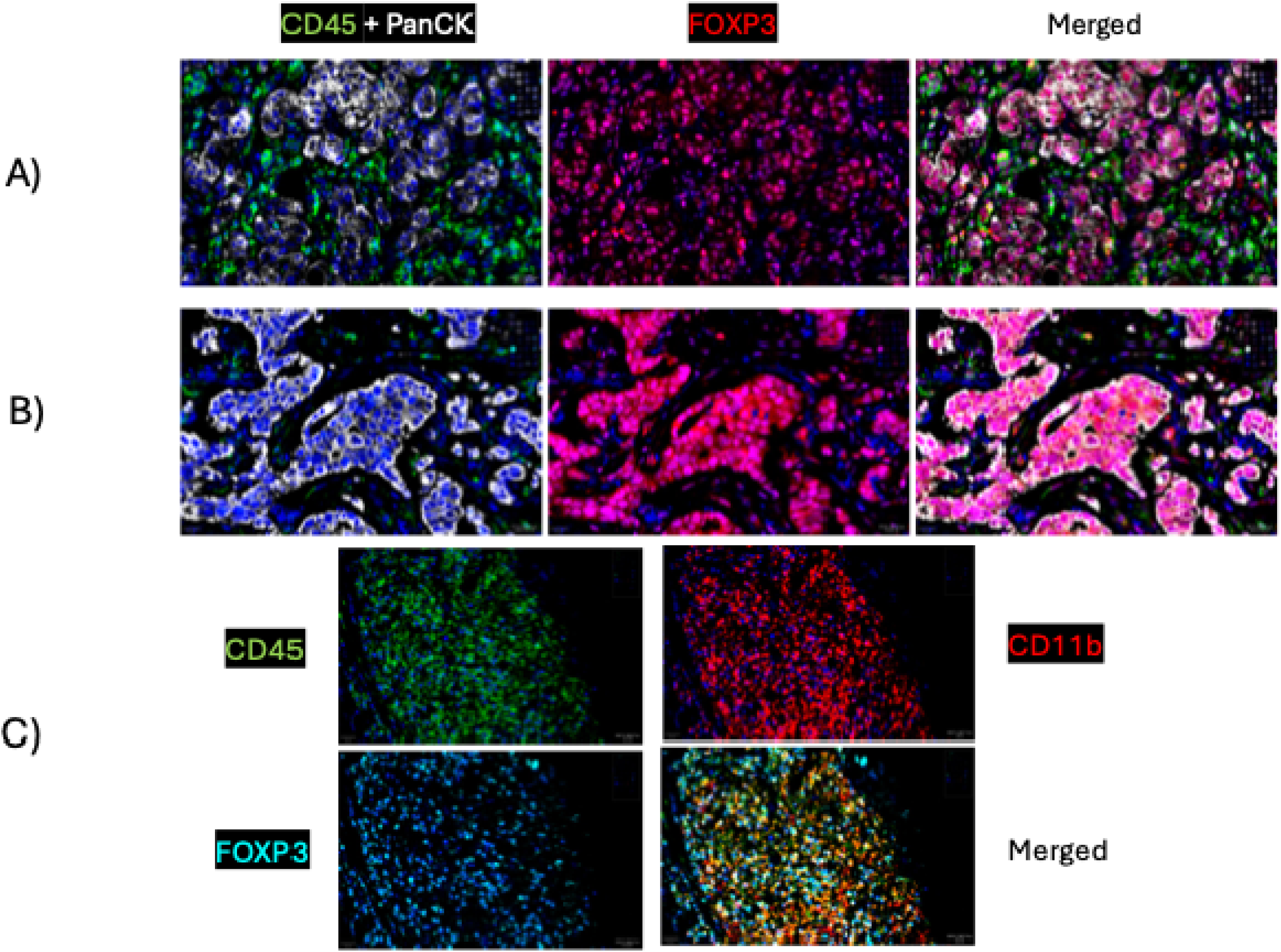
Non-canonical FoxP3 expression in tumor and myeloid populations. A-B) Representative multiplex immunofluorescence images from two independent tissue samples demonstrating FOXP3 expression within PanCK^+^ tumor niches. Individual and merged channels display CD45 (green), PanCK (white), FoxP3 (red), and DAPI (blue). The overlap of FoxP3 and PanCK signals confirms non-canonical FoxP3 expression within tumor cells. C) Representative multiplex immunofluorescence images demonstrating FOXP3 expression within the myeloid compartment. Individual and merged channels display CD45 (green), CD11b (red), FoxP3 (blue), and DAPI (blue). The merged panel demonstrates distinct co-localization of FoxP3 with CD45^+^/CD11b^+^ cells, identifying a population of FoxP3^+^ myeloid cells.

**Supplemental Figure 6:**
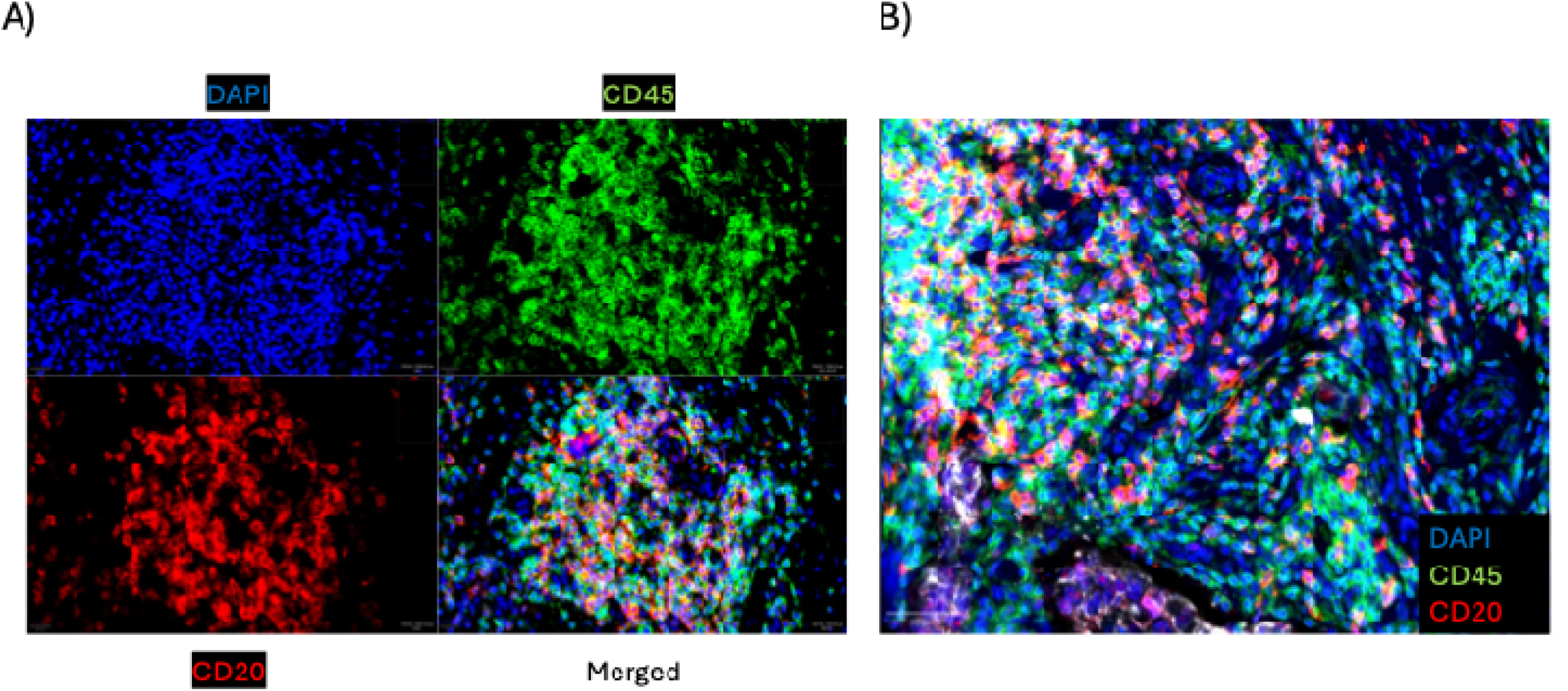
CD20+ B-cell staining pattern in tonsil and TNBC tissue. **(A)** Representative multiplex immunofluorescence images of tonsil tissue showing the expected staining pattern of CD20-positive B cells. Individual panels display DAPI (blue), CD45 (green), and CD20 (red), with the merged image shown at bottom right. In tonsil tissue, CD20 signal co-localizes with CD45-positive immune cells, consistent with canonical B-cell staining. **(B)** Representative multiplex immunofluorescence image of TNBC tissue showing CD20 staining.

**Supplemental Table 1:** CONCH-derived feature representations outperform other image encoders across all TME phenotyping tasks.

| Task |  | Test set AUROC |  |  |
| --- | --- | --- | --- | --- |
|  |  | *ResNet50* | *PLIP* | *CONCH* |
| **Immune** | *B-cell proliferation* | 0.6974 | 0.7755 | 0.8039 |
|  | *T-cell mediated cytotoxicity* | 0.7149 | 0.7703 | 0.7904 |
|  | *FoxP3-mediated immunosuppression* | 0.7996 | 0.8458 | 0.8469 |
|  | *Antigen processing and presentation* | 0.7450 | 0.7806 | 0.8260 |
| **Tumor** | *Angiogenesis* | 0.7042 | 0.7435 | 0.7605 |
|  | *Epithelial-to-mesenchymal transition* | 0.8110 | 0.7848 | 0.8120 |
|  | *Cell cycling* | 0.7768 | 0.7939 | 0.8100 |
| **Metabolic** | *Fatty acid metabolism* | 0.6323 | 0.6030 | 0.6649 |
|  | *Glycolysis* | 0.7996 | 0.8118 | 0.8276 |
|  | *Oxidative phosphorylation* | 0.6894 | 0.7145 | 0.7194 |

**Supplemental Table 2:** Models trained solely on tabular clinico-genomic data show limited ability to quantify TME biology.

| Task |  | Test set AUROC: Median [95% confidence interval] |  |  |
| --- | --- | --- | --- | --- |
|  |  | *Random Forest* | *Support Vector Machine* | *TabPFN* |
| **Immune** | *B-cell proliferation* | 0.4531 [0.3775; 0.5362] | 0.5697 [0.4961; 0.6493] | 0.5469 [0.4732; 0.6229] |
|  | *T-cell mediated cytotoxicity* | 0.5218 [0.4503; 0.6050] | 0.6154 [0.5484; 0.6911] | 0.6059 [0.5354; 0.6832] |
|  | *FoxP3-mediated immunosuppression* | 0.6266 [0.5542; 0.7069] | 0.7184 [0.6482; 0.7860] | 0.6922 [0.6201; 0.7633] |
|  | *Antigen processing and presentation* | 0.6018 [0.5241; 0.6724] | 0.6071 [0.5360; 0.6798] | 0.6012 [0.5278; 0.6766] |
| **Tumor** | *Angiogenesis* | 0.6031 [0.5290; 0.6780] | 0.5734 [0.4929; 0.6454] | 0.6156 [0.5472; 0.6839] |
|  | *Epithelial-to-mesenchymal transition* | 0.6294 [0.5581; 0.7008] | 0.6230 [0.5472; 0.6930] | 0.6501 [0.5802; 0.7210] |
|  | *Cell cycling* | 0.7509 [0.6853; 0.8099] | 0.8009 [0.7358; 0.8557] | 0.7900 [0.7269; 0.8458] |
| **Metabolic** | *Fatty acid metabolism* | 0.5984 [0.5278; 0.6755] | 0.5948 [0.5183; 0.6689] | 0.6053 [0.5310; 0.6808] |
|  | *Glycolysis* | 0.7179 [0.6467; 0.7854] | 0.7197 [0.6480; 0.7828] | 0.7274 [0.6572; 0.7899] |
|  | *Oxidative phosphorylation* | 0.5958 [0.5258; 0.6723] | 0.5954 [0.5181; 0.6709] | 0.6022 [0.5294; 0.6723] |

**Supplemental Table 3:**
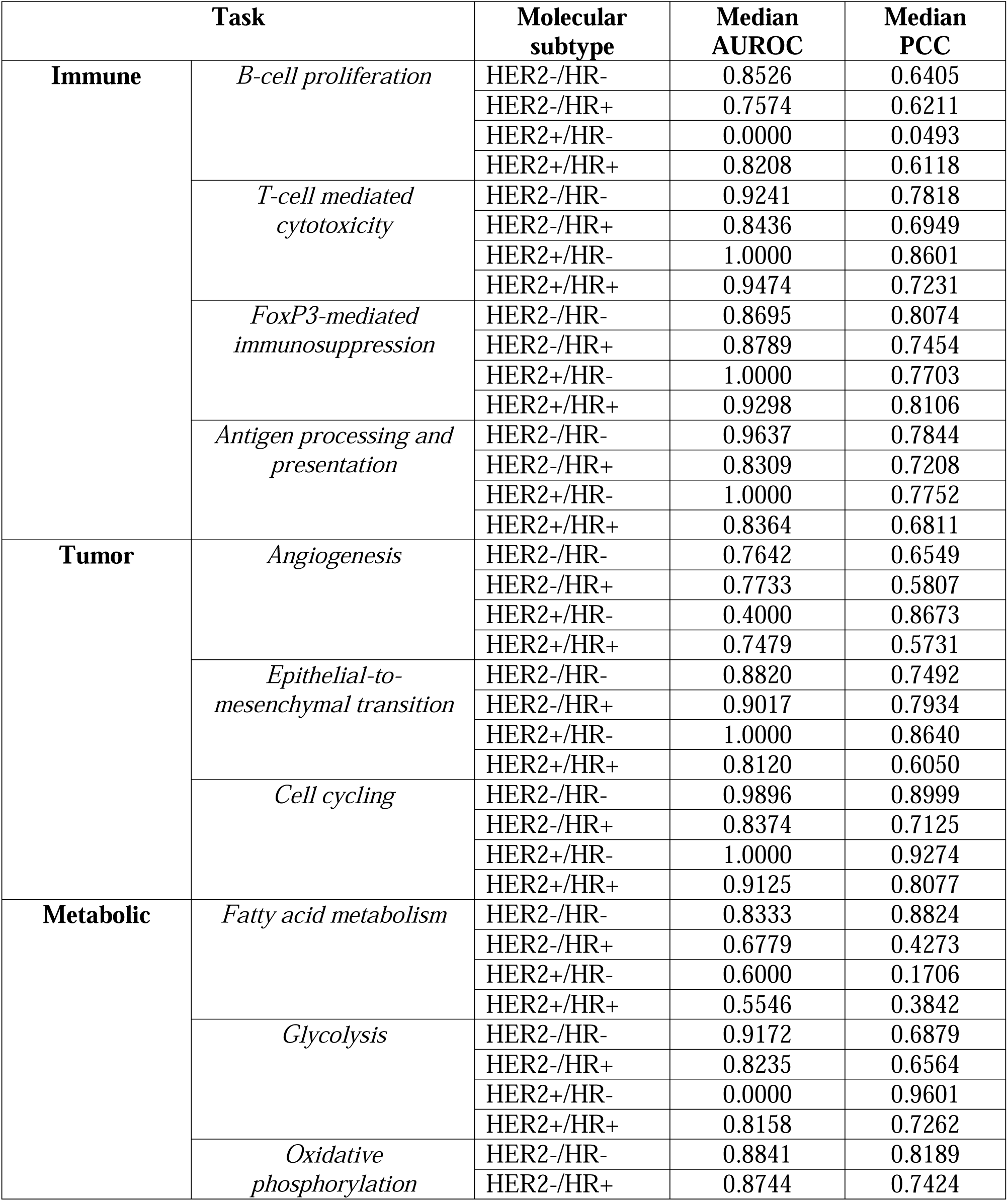
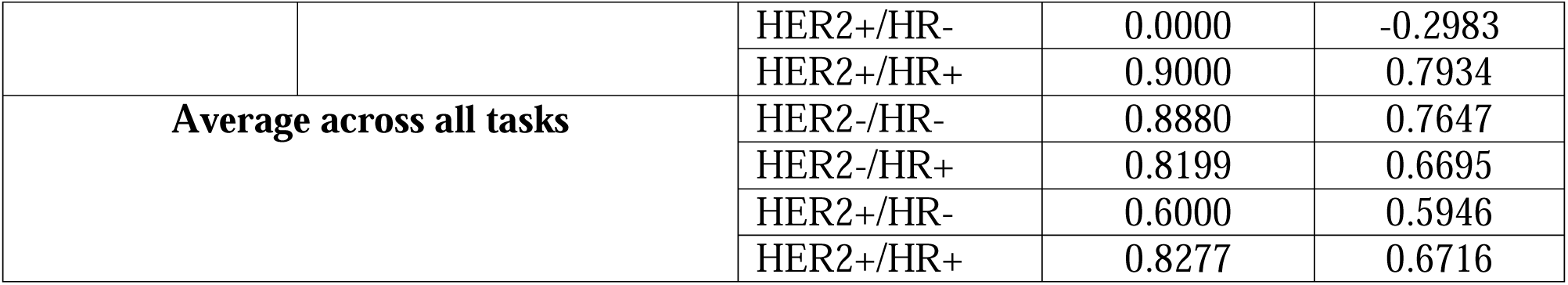
Performance of H&E WSI-based models vary across breast cancer molecular subtypes.

| Task |  | Molecular subtype | Median AUROC | Median PCC |
| --- | --- | --- | --- | --- |
| Immune | *B-cell proliferation* | HER2-/HR- | 0.8526 | 0.6405 |
|  |  | HER2-/HR+ | 0.7574 | 0.6211 |
|  |  | HER2+/HR- | 0.0000 | 0.0493 |
|  |  | HER2+/HR+ | 0.8208 | 0.6118 |
|  | *T-cell mediated cytotoxicity* | HER2-/HR- | 0.9241 | 0.7818 |
|  |  | HER2-/HR+ | 0.8436 | 0.6949 |
|  |  | HER2+/HR- | 1.0000 | 0.8601 |
|  |  | HER2+/HR+ | 0.9474 | 0.7231 |
|  | *FoxP3-mediated immunosuppression* | HER2-/HR- | 0.8695 | 0.8074 |
|  |  | HER2-/HR+ | 0.8789 | 0.7454 |
|  |  | HER2+/HR- | 1.0000 | 0.7703 |
|  |  | HER2+/HR+ | 0.9298 | 0.8106 |
|  | *Antigen processing and presentation* | HER2-/HR- | 0.9637 | 0.7844 |
|  |  | HER2-/HR+ | 0.8309 | 0.7208 |
|  |  | HER2+/HR- | 1.0000 | 0.7752 |
|  |  | HER2+/HR+ | 0.8364 | 0.6811 |
| Tumor | *Angiogenesis* | HER2-/HR- | 0.7642 | 0.6549 |
|  |  | HER2-/HR+ | 0.7733 | 0.5807 |
|  |  | HER2+/HR- | 0.4000 | 0.8673 |
|  |  | HER2+/HR+ | 0.7479 | 0.5731 |
|  | *Epithelial-to-mesenchymal transition* | HER2-/HR- | 0.8820 | 0.7492 |
|  |  | HER2-/HR+ | 0.9017 | 0.7934 |
|  |  | HER2+/HR- | 1.0000 | 0.8640 |
|  |  | HER2+/HR+ | 0.8120 | 0.6050 |
|  | *Cell cycling* | HER2-/HR- | 0.9896 | 0.8999 |
|  |  | HER2-/HR+ | 0.8374 | 0.7125 |
|  |  | HER2+/HR- | 1.0000 | 0.9274 |
|  |  | HER2+/HR+ | 0.9125 | 0.8077 |
| Metabolic | *Fatty acid metabolism* | HER2-/HR- | 0.8333 | 0.8824 |
|  |  | HER2-/HR+ | 0.6779 | 0.4273 |
|  |  | HER2+/HR- | 0.6000 | 0.1706 |
|  |  | HER2+/HR+ | 0.5546 | 0.3842 |
|  | *Glycolysis* | HER2-/HR- | 0.9172 | 0.6879 |
|  |  | HER2-/HR+ | 0.8235 | 0.6564 |
|  |  | HER2+/HR- | 0.0000 | 0.9601 |
|  |  | HER2+/HR+ | 0.8158 | 0.7262 |
|  | *Oxidative phosphorylation* | HER2-/HR- | 0.8841 | 0.8189 |
|  |  | HER2-/HR+ | 0.8744 | 0.7424 |
|  |  | HER2+/HR- | 0.0000 | -0.2983 |
|  |  | HER2+/HR+ | 0.9000 | 0.7934 |
| Average across all tasks |  | HER2-/HR- | 0.8880 | 0.7647 |
|  |  | HER2-/HR+ | 0.8199 | 0.6695 |
|  |  | HER2+/HR- | 0.6000 | 0.5946 |
|  |  | HER2+/HR+ | 0.8277 | 0.6716 |

**Supplemental Table 4:**
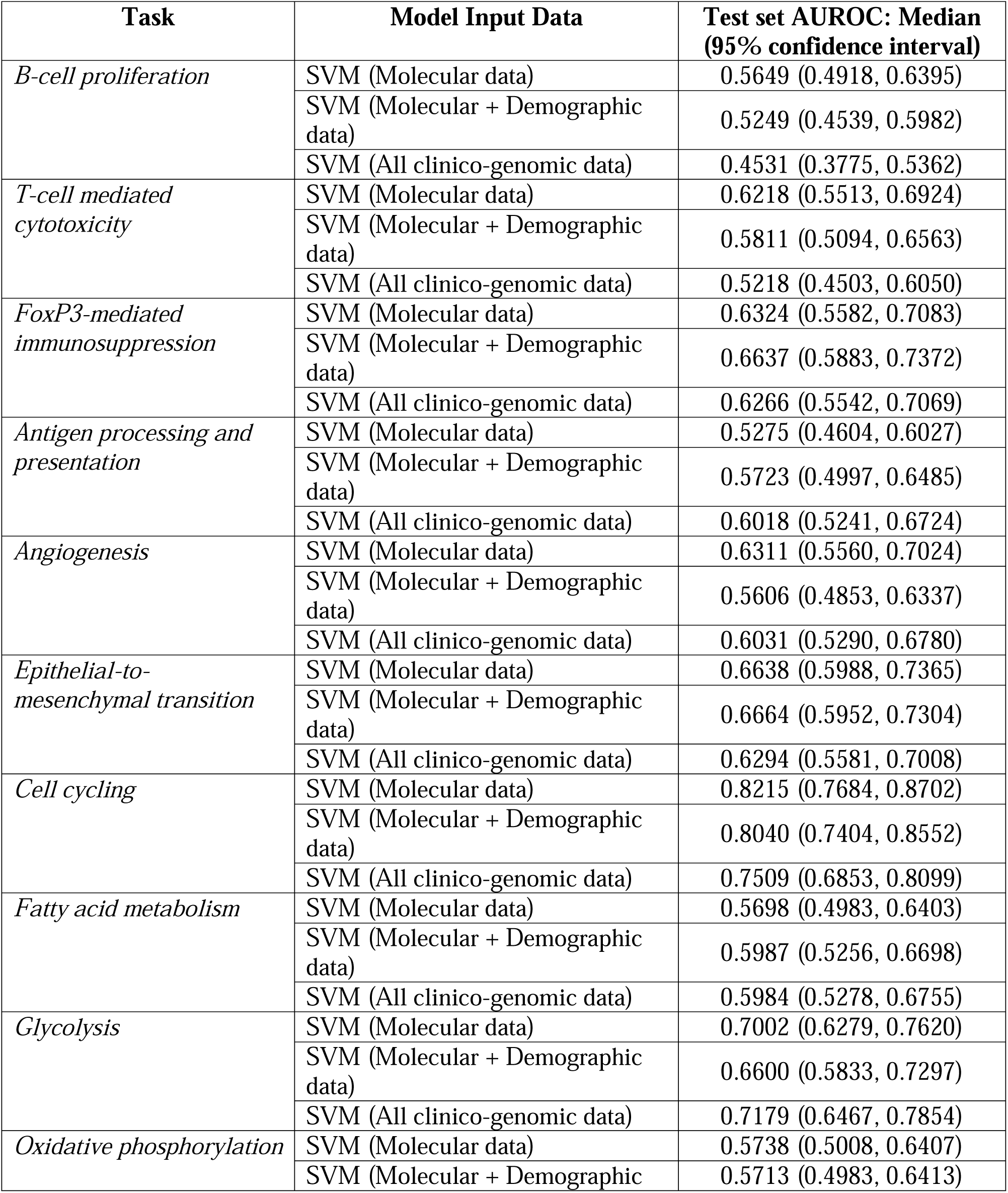
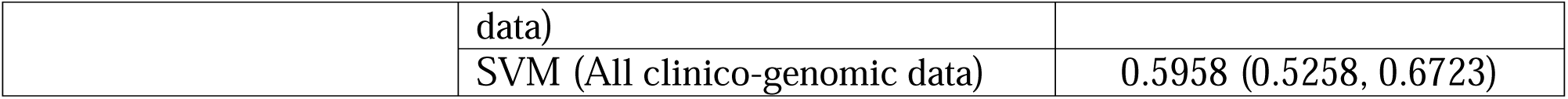
Ablation analyses indicate that predictive signal for assessing TME phenotypes in tabular clinico-genomic models is driven primarily by molecular data.

**Supplemental Table 5:**
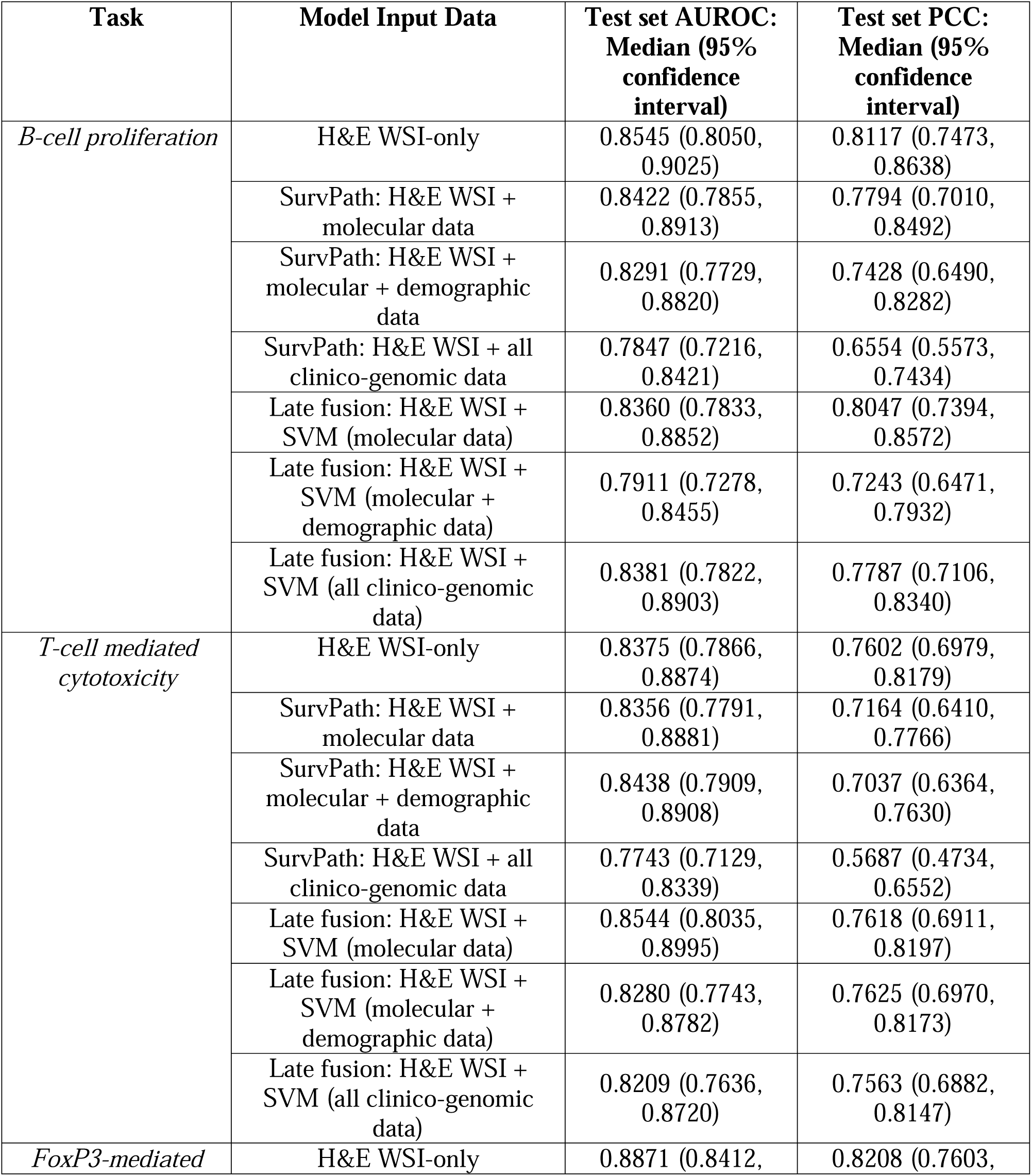
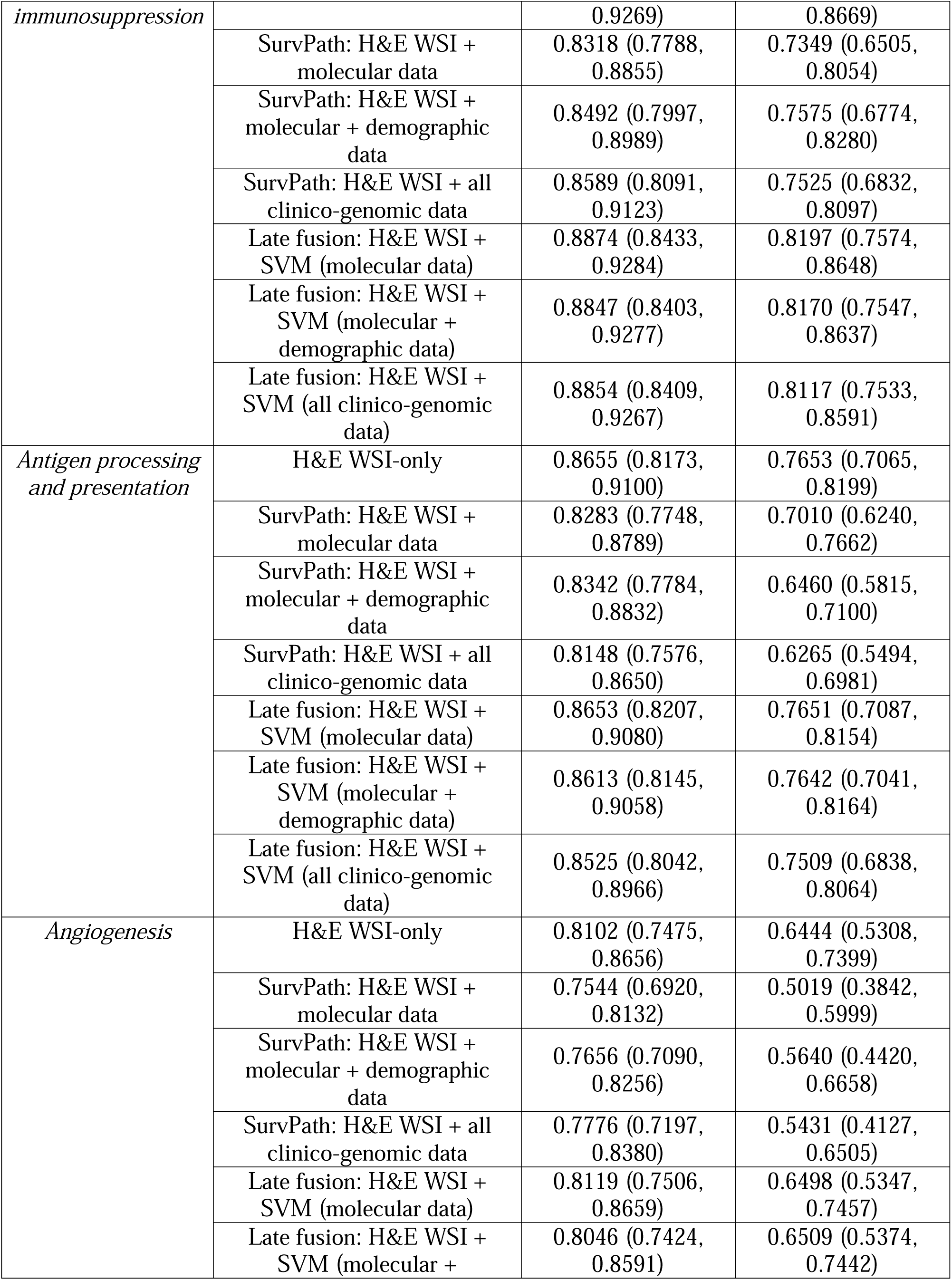
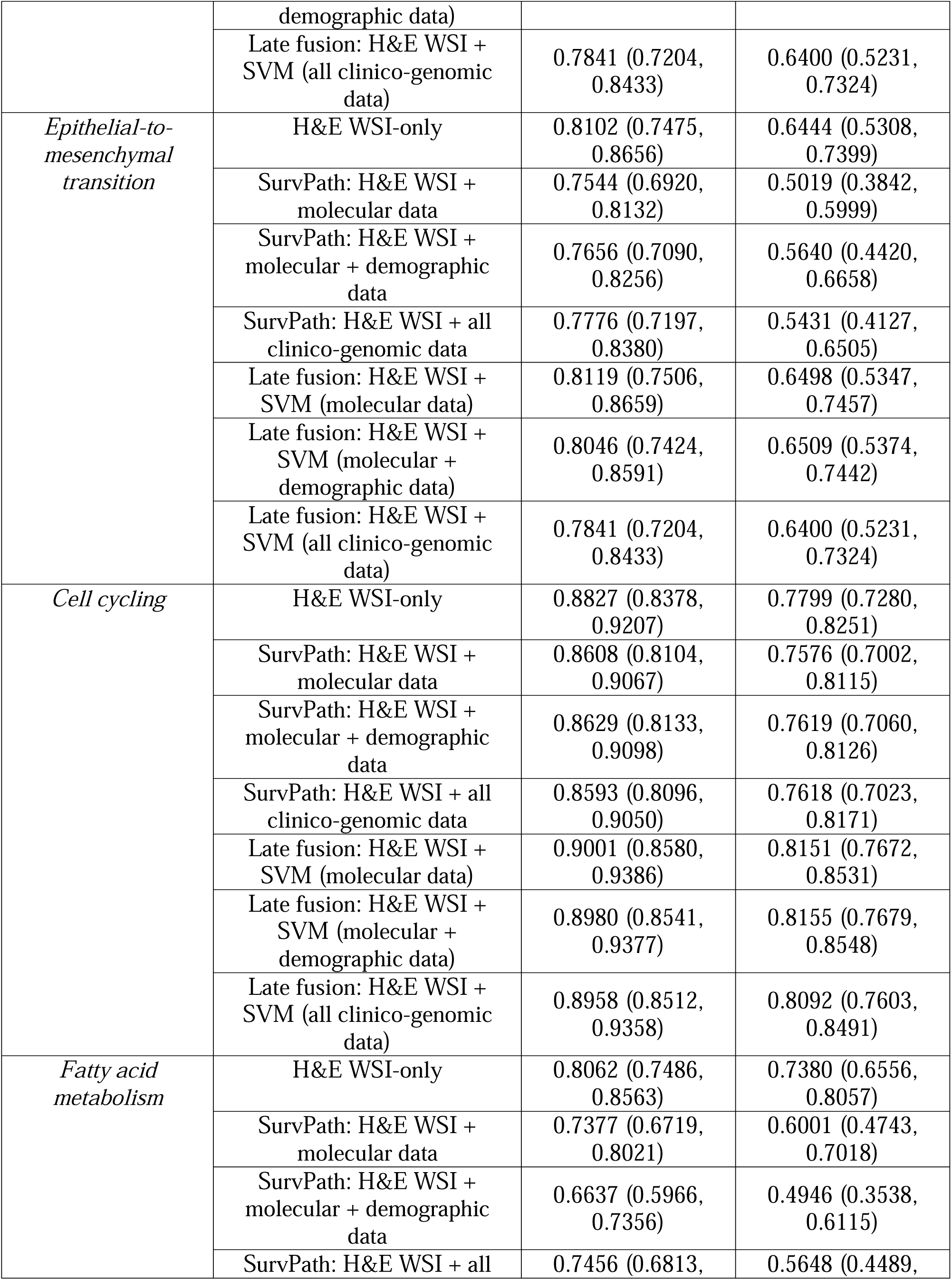
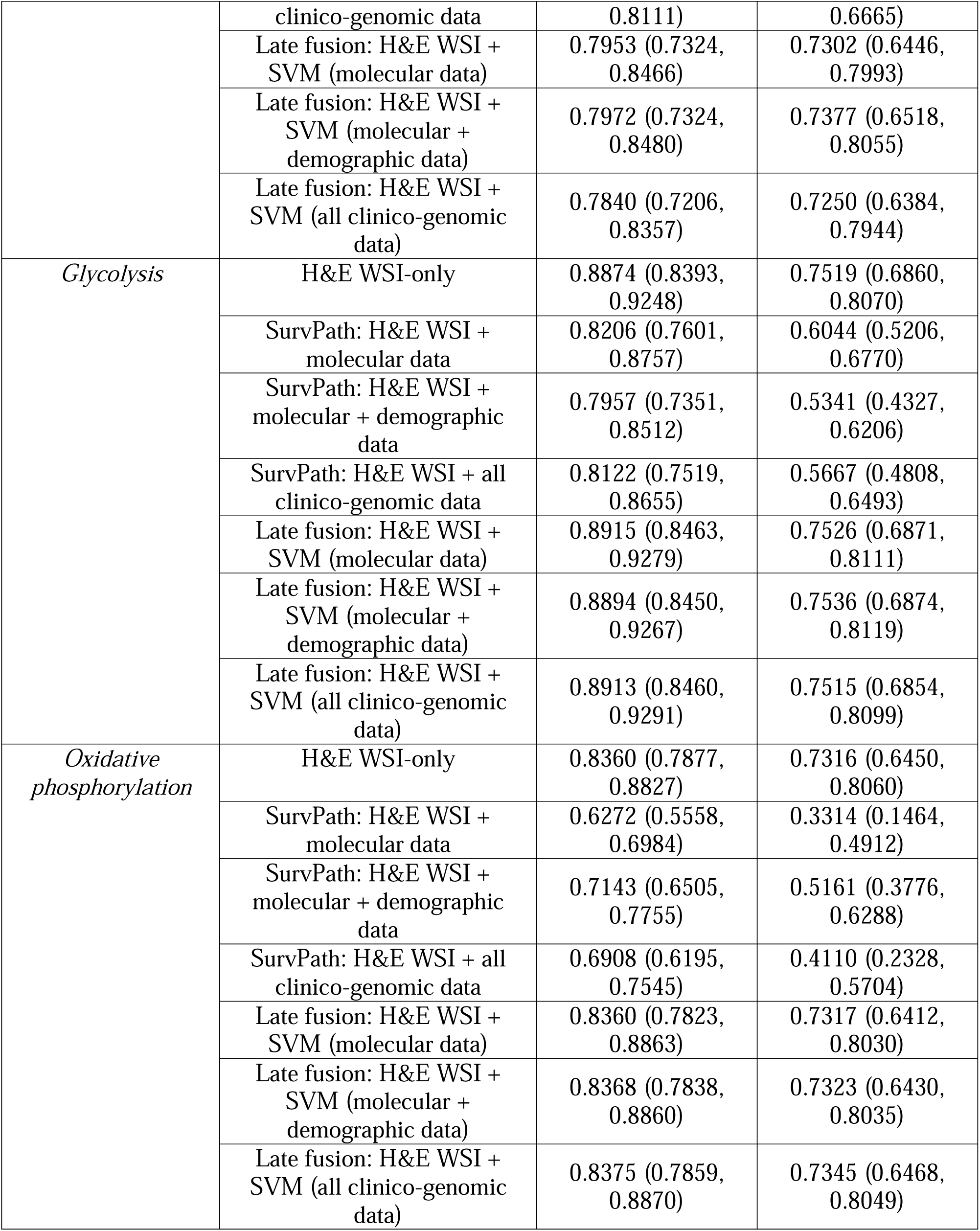
Ablation analyses suggest that adding tabular clinico-genomic data to H&E WSI models does not improve, and may reduce, model performance in assessing TME phenotypes.

## Notes

### Summary of Updates

reframing of manuscript to highlight identification of new biology; no new experiments.

## REFERENCES

1. Son, B. et al. The role of tumor microenvironment in therapeutic resistance. Oncotarget 8, 3933–3945 (2017).

2. de Visser, K. E. & Joyce, J. A. The evolving tumor microenvironment: From cancer initiation to metastatic outgrowth. Cancer Cell 41, 374–403 (2023).

3. Hemker, K., et al. Towards Spatial Transcriptomics-driven Pathology Foundation Models. https://arxiv.org/pdf/2602.14177v1 (2026).

4. Chen, W. et al. A visual–omics foundation model to bridge histopathology with spatial transcriptomics. Nature Methods 2025 22:7 22, 1568–1582 (2025).

5. Pizurica, M. et al. Digital profiling of gene expression from histology images with linearized attention. Nature Communications 2024 15:1 15, 9886- (2024).

6. Tsai, P. C. et al. Histopathology images predict multi-omics aberrations and prognoses in colorectal cancer patients. Nat. Commun. 14, (2023).

7. Chen, R. J. et al. Pan-cancer integrative histology-genomic analysis via multimodal deep learning. Cancer Cell 40, 865–878.e6 (2022).

8. Khatri, P., Sirota, M. & Butte, A. J. Ten years of pathway analysis: current approaches and outstanding challenges. PLoS Comput. Biol. 8, (2012).

9. Subramanian, A. et al. Gene set enrichment analysis: A knowledge-based approach for interpreting genome-wide expression profiles. Proc. Natl. Acad. Sci. U. S. A. 102, 15545– 15550 (2005).

10. Harbeck, N. et al. Breast cancer. Nature Reviews Disease Primers 2019 5:1 5, 66- (2019).

11. Shen, L. et al. Metabolic reprogramming in triple-negative breast cancer through Myc suppression of TXNIP. Proc. Natl. Acad. Sci. U. S. A. 112, 5425–5430 (2015).

12. Gonçalves, T., et al. Deep Learning-based Prediction of Breast Cancer Tumor and Immune Phenotypes from Histopathology. https://arxiv.org/abs/2404.16397v1 (2024).

13. Jaume, G. et al. Modeling Dense Multimodal Interactions Between Biological Pathways and Histology for Survival Prediction. Proceedings of the IEEE Computer Society Conference on Computer Vision and Pattern Recognition 11579–11590 (2023) doi:10.1109/CVPR52733.2024.01100.

14. Kim, J. et al. Limitations of Deep Learning Attention Mechanisms in Clinical Research: Empirical Case Study Based on the Korean Diabetic Disease Setting. J. Med. Internet Res. 22, e18418 (2020).

15. Wang, C. & Zhou, Y. Rethinking the role of attention mechanism: a causality perspective. Applied Intelligence 2024 54:2 54, 1862–1878 (2024).

16. Das, S. & Johnson, D. B. Immune-related adverse events and anti-tumor efficacy of immune checkpoint inhibitors. Journal for ImmunoTherapy of Cancer 2019 7:1 7, 306- (2019).

17. Dall’Olio, F. G. et al. Tumour burden and efficacy of immune-checkpoint inhibitors. Nature Reviews Clinical Oncology 2021 19:2 19, 75–90 (2021).

18. Stover, D. G. et al. The Role of Proliferation in Determining Response to Neoadjuvant Chemotherapy in Breast Cancer: A Gene Expression-Based Meta-Analysis. Clin. Cancer Res. 22, 6039–6050 (2016).

19. Filho, O. M. et al. Association of Immunophenotype With Pathologic Complete Response to Neoadjuvant Chemotherapy for Triple-Negative Breast Cancer: A Secondary Analysis of the BrighTNess Phase 3 Randomized Clinical Trial. JAMA Oncol. 7, 603–608 (2021).

20. Gao, G., Wang, Z., Qu, X. & Zhang, Z. Prognostic value of tumor-infiltrating lymphocytes in patients with triple-negative breast cancer: a systematic review and meta-analysis. BMC Cancer 20, (2020).

21. Drew, Y., Zenke, F. T. & Curtin, N. J. DNA damage response inhibitors in cancer therapy: lessons from the past, current status and future implications. Nature Reviews Drug Discovery 2024 24:1 24, 19–39 (2024).

22. Lord, C. J. & Ashworth, A. The DNA damage response and cancer therapy. Nature 2012 481:7381 481, 287–294 (2012).

23. Xiang, J., et al. A Multimodal Foundation Model of Spatial Transcriptomics and Histology for Biological Discovery and Clinical Prediction. https://arxiv.org/pdf/2604.03630v1 (2026).

24. Valanarasu, J. M. J. et al. Multimodal AI generates virtual population for tumor microenvironment modeling. Cell 189, 386–400.e19 (2026).

25. Redekop, E., et al. SPADE: Spatial Transcriptomics and Pathology Alignment Using a Mixture of Data Experts for an Expressive Latent Space. https://arxiv.org/pdf/2506.21857v1 (2025).

26. Jaume, G. et al. HEST-1k: A Dataset for Spatial Transcriptomics and Histology Image Analysis. Adv. Neural Inf. Process. Syst. 37, (2024).

27. Longo, S. K., Guo, M. G., Ji, A. L. & Khavari, P. A. Integrating single-cell and spatial transcriptomics to elucidate intercellular tissue dynamics. Nature Reviews Genetics 2021 22:10 22, 627–644 (2021).

28. Dosovitskiy, A., et al. An Image is Worth 16x16 Words: Transformers for Image Recognition at Scale. ICLR 2021 - 9th International Conference on Learning Representations https://arxiv.org/pdf/2010.11929 (2020).

29. Kaya, A., Bilik, I. & Stainvas, I. A Comparative Study of Vision Transformers and CNNs for Few-Shot Rigid Transformation and Fundamental Matrix Estimation. https://arxiv.org/pdf/2510.04794v1 (2025).

30. Hudson, T. J. et al. International network of cancer genome projects. Nature 464, 993–998 (2010).

31. Cerami, E. et al. The cBio Cancer Genomics Portal: An Open Platform for Exploring Multidimensional Cancer Genomics Data. Cancer Discov. 2, 401 (2012).

32. Balakrishnan, R., Harris, M. A., Huntley, R., Van Auken, K. & Michael Cherry, J. A guide to best practices for Gene Ontology (GO) manual annotation. Database 2013, (2013).

33. Nam, A. S., Chaligne, R. & Landau, D. A. Integrating genetic and non-genetic determinants of cancer evolution by single-cell multi-omics. Nat. Rev. Genet. 22, 3–18 (2021).

34. Subramanian, A. et al. Gene set enrichment analysis: a knowledge-based approach for interpreting genome-wide expression profiles. Proc. Natl. Acad. Sci. U. S. A. 102, 15545– 15550 (2005).

35. Barbie, D. A. et al. Systematic RNA interference reveals that oncogenic KRAS-driven cancers require TBK1. Nature 2009 462:7269 462, 108–112 (2009).

36. Schmid, P. et al. Pembrolizumab for Early Triple-Negative Breast Cancer. New England Journal of Medicine 382, 810–821 (2020).

37. Citron, M. L. et al. Randomized trial of dose-dense versus conventionally scheduled and sequential versus concurrent combination chemotherapy as postoperative adjuvant treatment of node-positive primary breast cancer: first report of Intergroup Trial C9741/Cancer and Leukemi…. J. Clin. Oncol. 21, 1431–1439 (2003).

38. Kather, J. N. et al. Deep learning can predict microsatellite instability directly from histology in gastrointestinal cancer. Nat. Med. 25, 1054–1056 (2019).

39. Lipkova, J. et al. Artificial intelligence for multimodal data integration in oncology. Cancer Cell 40, 1095–1110 (2022).

40. Lu, M. Y. et al. Data-efficient and weakly supervised computational pathology on whole-slide images. Nature Biomedical Engineering 2021 5:6 5, 555–570 (2021).

41. Lu, M. Y. et al. A visual-language foundation model for computational pathology. Nature Medicine 2024 30:3 30, 863–874 (2024).

42. Janowczyk, A., Zuo, R., Gilmore, H., Feldman, M. & Madabhushi, A. HistoQC: An Open-Source Quality Control Tool for Digital Pathology Slides. *JCO Clin*. Cancer Inform. 3, 1–7 (2019).

43. Faryna, K., van der Laak, J. & Litjens, G. Automatic data augmentation to improve generalization of deep learning in H&E stained histopathology. Comput. Biol. Med. 170, 108018 (2024).

44. Huang, Z., Bianchi, F., Yuksekgonul, M., Montine, T. J. & Zou, J. A visual–language foundation model for pathology image analysis using medical Twitter. Nature Medicine 2023 29:9 29, 2307–2316 (2023).

45. He, K., Zhang, X., Ren, S. & Sun, J. Deep Residual Learning for Image Recognition. Proceedings of the IEEE Computer Society Conference on Computer Vision and Pattern Recognition 2016-December, 770–778 (2015).

46. Shao, Z. et al. TransMIL: Transformer based Correlated Multiple Instance Learning for Whole Slide Image Classification. Adv. Neural Inf. Process. Syst. 3, 2136–2147 (2021).

47. Hollmann, N., Müller, S., Eggensperger, K. & Hutter, F. TabPFN: A Transformer That Solves Small Tabular Classification Problems in a Second. 11th International Conference on Learning Representations, ICLR 2023 https://arxiv.org/pdf/2207.01848 (2022).

48. Black, S. et al. CODEX multiplexed tissue imaging with DNA-conjugated antibodies. Nat. Protoc. 16, 3802–3835 (2021).

49. Tan, Y. et al. SPACEc: a streamlined, interactive Python workflow for multiplexed image processing and analysis. Nature Communications 2025 16:1 16, 10652- (2025).

50. Korsunsky, I. et al. Fast, sensitive and accurate integration of single-cell data with Harmony. Nat. Methods 16, 1289–1296 (2019).

